# A Sulfonated Thermoresponsive Injectable Gel for Sequential Release of Therapeutic Proteins to Protect Cardiac Function After a Myocardial Infarction

**DOI:** 10.1101/2022.02.07.479463

**Authors:** Adam J. Rocker, Maria Cavasin, Noah Johnson, Robin Shandas, Daewon Park

## Abstract

Myocardial infarction causes cardiomyocyte death and persistent inflammatory responses, which generates adverse pathological remodeling. Delivering therapeutic proteins from injectable materials in a controlled release manner may present an effective biomedical approach for treating this disease. A thermoresponsive injectable gel composed of chitosan, conjugated with poly(N-isopropylacrylamide) and sulfonate groups, was developed for spatiotemporal protein delivery to protect cardiac function after a myocardial infarction. The thermoresponsive gel delivered VEGF, IL-10 and PDGF in a sequential and sustained manner *in vitro*. An acute myocardial infarction mouse model was used to evaluate polymer biocompatibility and to determine therapeutic effects from the delivery system on cardiac function. Immunohistochemistry showed biocompatibility of the hydrogel, while the controlled delivery of the proteins reduced macrophage infiltration and increased vascularization. Echocardiography showed an improvement in ejection fraction and fractional shortening after injecting the thermal gel and proteins. A factorial design of experiments study was implemented to optimize the delivery system for the best combination and doses of proteins for further increasing stable vascularization and reducing inflammation using a subcutaneous injection mouse model. The results showed that VEGF, IL-10, and FGF-2 demonstrated significant contributions towards promoting long-term vascularization, while PDGF’s effect was minimal.

## INTRODUCTION

Cardiovascular disease is the leading cause of death in the United States [1]. Coronary heart disease is a subset of cardiovascular disease that develops as plaque accumulates in the coronary arteries which can lead to a myocardial infarction (MI). This ischemic disease accounts for one in seven deaths with 805,000 new or recurrent coronary attacks each year [2]. After MI, prolonged ischemia causes cardiomyocyte cell death and activates local inflammatory responses leading to reduced contractile function and adverse pathological remodeling, where ventricular dilation and wall thinning ultimately result in heart failure. Current treatment options such as bypass graft surgery, angioplasty and pharmaceuticals prevent further damage but cannot restore heart function [3]. To compensate for the inadequate regeneration ability of the adult heart, there is a critical need to develop novel strategies for therapeutic regeneration.

Injectable biomaterials can be developed as multiple protein delivery vehicles for therapeutically targeting the various pathologies resulting from MI. These biomedical therapies should aim to induce cardiac tissue regeneration, promote cardiomyocyte survival, and reduce ventricular wall stress, thereby attenuating pathological remodeling and improving cardiac function. Intramyocardial biomaterial injections have shown efficacy for increasing heart function and vascularization, reducing infarct size and fibrosis, increasing progenitor cell recruitment, and reducing cardiomyocyte apoptosis [4–6]. Additionally, biomaterials can be chemically modified to electrostatically bind to bioactive molecules for localized and controlled delivery of therapeutic proteins [7]. This binding interaction improves the stability of angiogenic proteins, reducing issues associated with short protein half-lives and rapid diffusion from treatment sites [8]. Sustained delivery of angiogenic and immunomodulatory factors from biomaterials can enhance cardiac repair following MI by increasing perfusion to ischemic tissue and reducing persistent inflammation [9].

Angiogenesis is the growth of nascent blood vessels from existing vasculature networks. This process is initiated through the release of growth factors which signal for the migration and proliferation of endothelial cells to form new vessels [10]. Later, a maturation process occurs where pericytes are recruited by platelet-derived growth factor (PDGF) to cover endothelial cell tubes and stabilize the vessels [11]. Vascular endothelial growth factor (VEGF) and fibroblast growth factor-2 (FGF-2) have shown to be the most significant initiators and regulators of physiological angiogenesis during growth, healing and in response to hypoxia [12]. However, when presented alone, VEGF delivery can cause leaky and immature vessel formation with poor function, and these vessels may regress quickly [13,14]. Conversely, late-stage angiogenic proteins like PDGF can interfere with early-stage factors when presented simultaneously, reducing vascularization [15–17]. Therefore, PDGF should be delivered sequentially, after VEGF or FGF2, to promote optimal therapeutic angiogenesis. To limit the damaging effects of the immune response, recent studies have shown that delivery of anti-inflammatory cytokines can optimize cardiac repair following MI in rodent models [18]. Interleukin-10 (IL-10) is a potent anti-inflammatory cytokine which acts on monocytes by deactivating them and suppressing many pro-inflammatory mediators [19,20]. Delivery of IL-10 in acute MI rodent models demonstrated a significant increase in left ventricular function and reduced infarct size [19,21]. While these results show promise, IL-10 has a short half-life *in vivo* and requires multiple injections for effective treatment, leading to undesirable side effects with a high treatment cost [22]. Similar to angiogenic proteins, IL-10 possesses strong heparin-binding affinity properties (K_d_ = 54 ± 7 nM) [23]. With IL-10’sstrong electrostatic affinity for the sulfonate groups on heparin, a highly sulfonated hydrogel delivery system may provide a localized and sustained release of IL-10, reducing issues associated with high therapeutic doses.

In order to address this therapeutic strategy, a novel, sequential protein delivery system composed of a glycol chitosan (GC) backbone with poly(N-isopropylacrylamide) (PNIPAM) side chains and conjugated with sulfonate groups (S-GC-PNIPAM) has been developed. The GC backbone can be easily functionalized as it contains different functional groups to provide specific chemical modifications. Temperature responsive hydrogels are characteristic of aqueous solutions solidifying into hydrogels when a critical temperature is reached, such as physiological temperature. PNIPAM is a polymer with a lower critical solution temperature of 32 °C and has been used to provide temperature responsive properties for the controlled release of drugs [24,25]. Although heparin has been shown to have beneficial effects in drug delivery, heparin is difficult to modify, is susceptible to loss of sulfonate groups, and suffers from batch-to-batch variation [26]. This has led to the development of heparin mimicking polymers that incorporate sulfonate groups. Creating a delivery system that can sequentially deliver VEGF first, followed by IL-10, and lastly PDGF, according to their binding affinity to sulfonate groups on heparin, may promote optimal cardiac repair after MI. The quick delivery of VEGF initiates the angiogenesis response, potentially protecting cardiomyocytes from necrosis [27]. The release of IL-10 may prevent undue inflammatory injury to the heart [28]. The final delivery of PDGF should stabilize new vessels with pericytes, providing substantial revascularization to hypoxic heart tissue [29]. In the present study, the efficacy of this delivery system was evaluated for the spatiotemporal and sequential delivery of VEGF, IL-10, and PDGF towards the protection of cardiac function following MI.

## MATERIALS AND METHODS

### Materials

N-hydroxysuccinimide (NHS), 1,3-propane sultone (PS), dimethyl sulfoxide (DMSO), glycol chitosan (GC) and bovine serum albumin (BSA) were purchased from Sigma Aldrich (St. Louis, MO, USA). 1-ethyl-3-(−3-dimethylaminopropyl) carbodiimide hydrochloride (EDC) was purchased from Fisher Scientific (Pittsburgh, PA, USA). 2-Bromo-2-methylpropionic acid (BMPA), 1,1,4,7,10,10-hexamethyltriethylenetetramine (HMTETA), and copper(I) bromide (CuBr) were purchased from Alfa Aesar. Goat anti-rabbit IgG secondary antibody Alexa Fluor 594, rabbit anti-goat IgG secondary antibody Alexa Fluor 594, goat anti-rat IgG secondary antibody Alexa Fluor 488, cluster of differentiation 31 (CD31, rat IgG2a) primary antibody and alpha-smooth muscle actin (α-SMA, rabbit IgG) primary antibody were purchased from Thermo Fisher Scientific (Waltham, MA, USA). Von Willebrand factor (VWF, sheep IgG) primary antibody and cluster of differentiation 68 (CD68, rabbit IgG) primary antibody were purchased from Abcam (Cambridge, Ma). Human PDGF-BB standard ABTS (enzyme-linked immunosorbent assay) ELISA development kit, murine IL-10 standard ABTS ELISA development kit, murine VEGF standard ABTS ELISA development kit, recombinant murine FGF-2, recombinant human PDGF-BB, recombinant murine PDGF-BB, recombinant murine IL-10 and recombinant murine VEGF165 were purchased from Peprotech (Rocky Hill, NJ, USA). 10 % formalin was purchased from JT Baker (Phillipsburg, NJ, USA). N-isopropylacrylamide (NIPAm) and sucrose (RNASE & DNASE free) was purchased from VWR Life Science (Radnor, PA, USA). Dapi flouromount-G was purchased from Electron Microscope Sciences (Hartfield, PA, USA). Spectra/Por dialysis membranes (MWCO: 3500–5000 and 12,000–14,000 Da) were purchased from Spectrum Laboratories (Rancho Dominguez, CA).

### Equipment

Proton nuclear magnetic resonance (^1^H NMR) was performed on a Varian Inova 500 NMR Spectrometer. Fourier transform infrared spectroscopy (FTIR) was performed on a Nicolet 6700 FTIR Spectrometer and samples were run on polyethylene infrared sample cards. Scanning electron microscope (SEM) images and elemental analysis measurements by energy-dispersive spectroscopy (EDS) were taken using a JEOL (Peabody, MA) JSAM-6010la. Rheological characterization was performed using a TA Instruments Discovery HR-2 rheometer. ELISA color development was monitored with an ELISA plate reader (BioTek Synergy 2 Multi-Mode Reader) at 405 nm with wavelength correction set at 650 nm. Echocardiographic measurements were obtained using FUJIFILM VisualSonics Vevo 2100 equipped with a 30 MHz transducer. Tissue was sectioned using a CryoStar NX70 Cryostat. Brightfield images were taken using a Nikon Eclipse Ti-E. Confocal images were taken using a Zeiss LSM 780 spectral microscope. Zeiss ZEN blue software was used to quantify different parameters for image analysis from confocal images.

### Synthesis of S-GC-PNIPAM

For the preparation of bromoisobutyryl-terminated glycol chitosan (GC), 2-bromo-2-methylpropionic acid (BMPA, 0.13 g), EDC (0.21 g) and NHS (0.13 g) were dissolved in 8 ml of ultrapure water. After the mixture was activated at room temperature for 2 h, a 1% solution of GC (0.21 g, ultrapure water) was added to the flask, and stirred for 24 h at room temperature. At the end of the reaction, the solution was dialyzed against ultrapure water (4 × 4 L) with dialysis membrane (MWCO, 3.5 kDa) at 4 °C for 3 days. The product was subsequently lyophilized at −45 °C for 48 h and recovered for further conjugation.

Glycol chitosan-PNIPAM (GC-PNIPAM) was synthesized from using bromoisobutylryl-terminated GC as an initiator for the atom transfer radical polymerization (ATRP) of PNIPAM. For the ATRP reaction, PNIPAM was synthesized using a molar feed ratio [NIPAM, 2.1 g)]:[GC-bromine (GC-Br, 0.13 g)]:[Copper bromide (CuBr, 13.2 mg)]:[HMTETA, 32.2 µl)] of 60:1:1:1.2, following typical conditions for ATRP. NIPAM, GC-Br, and HMTETA were first added to a flask containing 15 ml of ultrapure water. After NIPAM was completely dissolved, the reaction mixture was degassed by bubbling nitrogen through the solution for 30 min. Then, CuBr was added into the mixture under a nitrogen atmosphere. The reaction mixture was then purged with nitrogen for another 10 min. The flask was sealed under a nitrogen atmosphere. The polymerization was performed under continuous stirring at room temperature for 48 h. At the end of the reaction, the solution was dialyzed against ultrapure water (5 × 4 L) with dialysis membrane (MWCO, 3.5 kDa) for 5 days. The product was lyophilized at −45 °C for 48 h.

For the sulfonation reaction, GC-PNIPAM (0.1 g) and potassium tert-butoxide (0.012 g, 10% molar) were dissolved in 5 mL of ultrapure water under a nitrogen atmosphere. Propane sultone (0.611 g, 5 molar excess) was added slowly to the flask and the sulfonation reaction was performed for 3 days at room temperature under a nitrogen atmosphere. At the end of the reaction, the solution was dialyzed against ultrapure water (5 × 4 L) with dialysis membrane (MWCO, 3.5 kDa) for 5 days. The resultant product, sulfonated-GC-PNIPAM (S-GC-PNIPAM), was lyophilized at −45 °C for 48 h.

### Molecular structure characterization

Proton nuclear magnetic resonance (^1^H NMR) was used to confirm the molecular structure of GC, GC-Br and GC-PNIPAM after each reaction. Samples (3-5 mg) were dissolved in 600 µl D_2_O and analyzed with a Varian Inova 500 NMR Spectrometer. Spectra were processed using ACD 1D NMR Processor software (Advanced Chemistry Development, Inc.). GC-Br, GC-PNIPAM and S-GC-PNIPAM samples were examined with Fourier transform infrared (FTIR) spectroscopy to confirm the conjugation of the sulfonate groups. Samples were dissolved in DMSO and placed on polyethylene IR cards which were then analyzed by a Nicolet 6700 (Thermo Fisher Scientific).

### Elemental analysis and scaffold morphology

GC-PNIPAM (4%, saline) and S-GC-PNIPAM (4%, saline) solutions were gelled at 37 °C for 10 minutes. The gelled samples were flash frozen in liquid nitrogen and quickly cut in half to expose the internal structure of the scaffolds, followed by lyophilization to dry the samples. To determine the amount of sulfonation on S-GC-PNIPAM, elemental analysis was performed using EDS to analyze the inner scaffold surface. Scanning electron microscopy (SEM) was used to examine the scaffold structure and morphology on the micrometer scale. For morphology assessment by SEM, samples were sputter coated with gold and palladium.

### Thermal gelling properties

Rheological characterization of S-GC-PNIPAM (4 wt%, 100 µl) was performed from 25 °C to 45 °C at a ramp rate of 1 °C/minute with an 8 mm parallel plate, a frequency of 1 Hz and stress of 0.05 pascals to determine the gelation temperature of the scaffold.

### Protein release study

S-GC-PNIPAM (4 wt%, 50 µl PBS) solutions were left to dissolve overnight at 4 °C with samples prepared in triplicate. Samples were loaded with 100 ng each of VEGF, IL-10 and PDGF by adding the proteins and mixing with a pipettor. The gels were formed in 2 ml vials in a 37 °C incubator for 5 minutes to allow for gel stabilization, and then 1 ml of warm (37 °C) release solution (PBS) was added to the vials. After 5 minutes, 1 ml of release solution was removed from each sample for the first time point, and 1 ml of fresh release solution was added. The first time point was used to determine the loading efficiencies of each protein in the hydrogels. Samples were then taken every 24 hours for four weeks, with weekly samples taken thereafter for the remaining 3 weeks, and immediately placed in a −80 °C freezer for further analysis. A sandwich ELISA was used to quantify the concentration of proteins in the release solution samples eluted from the hydrogels. The protocol from the manufacturerwas followed for each ELISA kit.

### Acute MI permanent ligation mouse model

Animal procedures for the myocardial infarction permanent ligation study were approved by the Institutional Animal Care and Use Committee (IACUC). C57BL/6J mice (The Jackson Laboratory) weighing 24−28 g were maintained on a light/dark cycle with access to food and water ad libitum. The study involved 4 – 6 mice per injection group (saline, bolus VEGF, IL-10 and PDGF (bolus 3 proteins), S-GC-PNIPAM, S-GC-PNIPAM + VEGF, IL-10 and PDGF (S-GC-PNIPAM + 3 proteins)) that survived 28 days after the permanent ligation procedure. MI was induced by coronary artery ligation as described previously [30,31]. Briefly, a left thoracotomy was performed through the fourth intercostal space and the pericardium was opened to expose the heart. The left anterior descending artery (LAD) was isolated. 8-0 suture was used to permanently ligate the LAD approximately 2 mm distal from its origin. Infarction of the left ventricular wall was confirmed by the tissue color change from pink to pale after ligation. Intramyocardial injections were performed after confirmation of MI. Saline (30 μl), bolus 3 proteins (500 ng of each protein in 30 μl saline), S-GC-PNIPAM (1 wt%, 30 μl saline) or S-GC-PNIPAM + 3 proteins (1 wt%, 500 ng of each protein in 30 μl saline) was intramyocardially injected at three equidistant points (10 μl each injection) around the ligation site through a 31-gauge needle. A control sham group (n = 3) was used as a baseline comparison, and no injection was administered for this group. The chest cavity was closed by suturing the incision in the third or fourth intercostal space with absorbable suture.

### Evaluating cardiac function with echocardiography

Serial transthoracic echocardiography was performed 0, 7, 14 and 28 after surgery to measure left ventricular function as described previously [30]. Briefly, mice were anaesthetized and maintained on a heated platform. Standard echocardiography was performed using a FUJIFILM VisualSonics Vevo 2100 equipped with a 30 MHz linear ultrasonic transducer in a phased array format. Two-dimensional m-mode targeted recordings were obtained from short axis views of the left ventricle at the papillary muscle and functional measurements were averaged from three cardiac cycles.

### Cardiac tissue harvest

Hearts were isolated and harvested 28 days after myocardial infarction and injection as described previously [30]. Briefly, mice were anaesthetized, and the thoracic cavity was opened with the heart still beating. Potassium chloride (10 wt%, saline) was injected through the left ventricle to arrest the heart in diastole, after which the heart was removed. The hearts were rinsed with PBS, embedded in OCT, and frozen at −80 °C. Cardiac tissue was sectioned transversely at a thickness of 5 μm beginning at the apex.

### Optimization of protein delivery system with factorial design of experiments dosage study and subcutaneous injection mouse model

Animal procedures were approved by IACUC and performed as described previously [32]. This study involved 3 mice per injection group with 8 different treatment groups and a saline control group. Treatment groups were determined from a literature review on different protein dosages used for therapeutic angiogenesis studies in murine animal models. A two-level half fractional factorial study design was formulated to select the most effective combinations of proteins and doses at inducing therapeutic angiogenesis and reducing inflammation, while potentially promoting functional cardiac recovery after MI. The two-levels are in reference to choosing the upper and lower doses for each protein. Based on the literature analysis, 3 µg was chosen as the upper dose and 0.3 µg for the lower dose (one-tenth of upper dose). The half fractional design involves performing half of the total runs. This design allows us to estimate all main effects between the proteins, which will help determine the contribution of each combination and optimal dose. The standard design formula of 2^(k-p)^ was used, with k = 4 factors, and p = 1, representing the half fractional factorial design, resulting in 2^3^ = 8 dose groups. JMP Pro software was used to setup the study design for the different dose groups and produce the statistical models necessary to determine the ideal protein combinations (Table 1). Saline (30 μl) or S-GC-PNIPAM (1 wt%, 30 μl saline) loaded with varying doses of 4 different proteins (0.3 µg or 3 µg each: VEGF, IL-10, PDGF, and FGF-2; 8 groups) was injected subcutaneously in the lower back of the mice through a 27-gauge needle.

**Table 1.** Dosage groups according to two-level half fractional factorial study design.

| **Protein** | **VEGF** | **IL-10** | **PDGF** | **FGF2** | **Functional Vascularization** | **Mature Vascularization** | **Inflammation** |
| --- | --- | --- | --- | --- | --- | --- | --- |
| Dose group 1 | 3 μg | 3 μg | 3 μg | 3 μg | ? | ? | ? |
| Dose group 2 | 3 μg | 3 μg | 0.3 μg | 0.3 μg | | | |
| Dose group 3 | 3 μg | 0.3 μg | 3 μg | 0.3 μg | | | |
| Dose group 4 | 3 μg | 0.3 μg | 0.3 μg | 3 μg | | | |
| Dose group 5 | 0.3 μg | 3 μg | 3 μg | 0.3 μg | | | |
| Dose group 6 | 0.3 μg | 3 μg | 0.3 μg | 3 μg | | | |
| Dose group 7 | 0.3 μg | 0.3 μg | 3 μg | 3 μg | | | |
| Dose group 8 | 0.3 μg | 0.3 μg | 0.3 μg | 0.3 μg | | | |
| Saline |  |  |  |  |  |  |  |

Subcutaneous tissue was harvested from the area of the injection site 28 days after injection. Mice were euthanized and the subcutaneous tissue (2 cm × 2 cm) was collected. The tissue was fixed in formalin (10%, PBS) overnight, cryoprotected with 30% sucrose in PBS for 24 h, embedded in OCT compound, and frozen at −80 °C. The tissue was sectioned transversely with a thickness of 10 µm and collected on glass slides.

### Histology

For assessment of fibrotic tissue, Masson trichrome staining was used to stain collagen fibers in heart sections as described previously [30]. Infarct size was measured from brightfield images of five trichrome stained cardiac tissue sections (960 μm, 1440 μm, 1920 μm, 2400 μm, and 2880 μm away from the apex) and averaged.

### Immunohistochemistry

For evaluation of angiogenesis and inflammation, cells were stained and identified as described previously [30,32]. Sections were stained with primary antibodies CD31 (1:50), VWF (1:50), α-SMA (1:250), and/or CD68 (1:500) for endothelial cells, vascular endothelial cells, pericytes, and macrophages, respectively. The sections were stained with secondary antibodies Alexa Fluor 488 (1:500) for CD31 and/or the associated Alexa Flour 594 (1:500) for VWF, α-SMA, and CD68. DAPI Fluoromount-G was used to stain nuclei and mount the sections.

Confocal images of the immunohistochemically stained cardiac tissue were obtained from around the infarct zone of the injection site (approximately 1920 μm from the apex) with z-stack projections (4 μm thickness, 1 μm steps). Vessel counts were computed from six to eight randomly selected images of the infarct or peri-infarct area and averaged. DAPI-associated stains were used to designate cells positive for each specific stain.

Confocal images of the immunohistochemically stained subcutaneous tissue were obtained from around the injection site with z-stack projections (4 μm thickness, 1 μm steps). Cell counts were computed from five randomly selected images and averaged. DAPI-associated stains were used to designate cells positive for each specific stain.

### Statistical Analysis

All results are expressed as means ± standard deviation. JMP Pro software generated the different dosage groups for the factorial DOE study and analyzed the resulting data. Two-tailed t-test assuming unequal variances was used to determine significant differences between two groups. Analysis of variance (ANOVA) followed by Tukey’s post hoc test was used to determine significant differences between three or more groups. Statistical significance (*) was considered when p < 0.05.

## RESULTS AND DISCUSSION

### Synthesis of S-GC-PNIPAM and polymer characterization

The synthesized S-GC-PNIPAM was designed to mimic the electrostatic interaction between the sulfonate groups on heparin and positively charged proteins, providing similar biofunction to the proteoglycan, while having the advantage of being a consistently reproducible thermoresponsive gel. S-GC-PNIPAM contains amide and glycosidic bonds to provide sites for degradation. The initial step in the synthesis of S-GC-PNIPAM involves conjugating BMPA to the primary amine groups on the GC backbone to act as the initiator for the ATRP reaction of PNIPAM (Figure 1). The carboxylic acid groups on BMPA are first activated into reactive esters by utilizing EDC/NHS chemistry. The resultant CS-Br initiators are produced from the esters undergoing nucleophilic substitution reactions with the amine groups on GC, which forms stable amide linkages.

**Figure 1.**
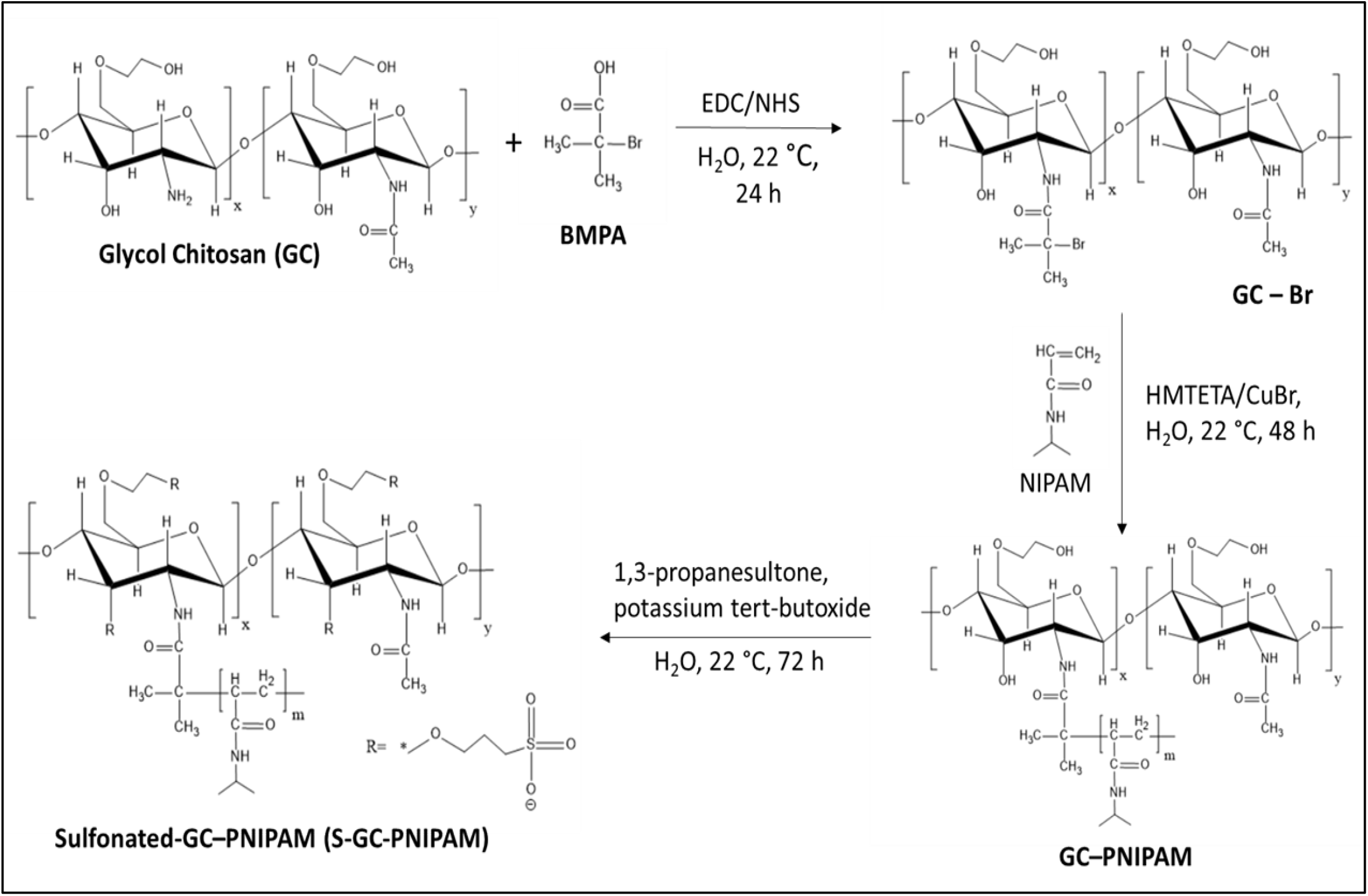
Reaction sequence of the sulfonated thermoresponsive gel S-GC-PNIPAM.

Using ^1^H NMR, the chemical structure of CS-Br was characterized. By comparison to the spectrum of GC (Figure S1A), a new chemical shift was identified at δ = 1.19 ppm. This shift corresponds to the methyl protons (d′, C(Br)-<u>CH</u>_<u>3</u>_) on the 2-bromoisobutyryl groups in the spectrum of CS-Br (Figure S1B), demonstrating that the ATRP initiators have been produced. Subsequently, PNIPAM side chains were synthesized via ATRP from NIPAM and CS-Br. Different lengths of PNIPAM can be synthesized by varying the ATRP time. The NMR spectrum for GC-PNIPAM (Figure S1C) exhibits chemical shifts at δ = 0.9 – 1.7 ppm which are attributable to the methylene protons (f, CH – <u>CH</u>_<u>2</u>_). The chemical shifts in the area of 1.8 – 2.7 ppm are associated with the methylidyne protons next to the carbonyl group (e, <u>CH</u> – C = O) in the PNIPAM chain, and the methyl protons (d, O = C(NH) – <u>CH</u>_<u>3</u>_ of GC; d′′, NH – CH – <u>CH</u>_<u>3</u>_ of PNIPAM).

FTIR was used to confirm the conjugation of the sulfonate groups. GC-Br and GC-PNIPAM provided baseline spectra to compare with S-GC-PNIPAM to identify the sulfonate peak. The full spectrum for the polymers shows a broad peak between the frequency range of 3500 – 3200, corresponding to the stretching of the O-H bond on primary alcohols. There is a reduction in the size of this peak for S-GC-PNIPAM compared to the other polymers, likely confirming the sulfonation of the free hydroxyls on the GC backbone (Figure S2A). Peaks appearing from the sulfonated polymer spectrum represent changes in bonds due to the sulfonation process, and these specific IR shifts occur between the frequency range of 1050 – 1025 cm^-1^. The IR spectrum for S-GC-PNIPAM confirms the presence of the sulfonate groups as this shift does not appear in GC-BR or GC-PNIPAM spectra (Figure S2B).

The incorporation of PNIPAM into hydrogels has been heavily researched due to the polymer’s reversible thermal gelling properties. This polymer exhibits a transition from a soluble liquid solution to a solid gel once the lower critical solution temperature (LCST) threshold has been crossed, or when injected at body temperature (37 °C), trapping any proteins in the gel during this transition. PNIPAM is composed of hydrophilic and hydrophobic segments which provide these thermoresponsive properties. The hydrophilic interactions dominate at room temperature allowing for PNIPAM solubility in water. At temperatures above the LCST, the hydrophobic interactions become dominate and the aqueous PNIPAM solution turns into a gel. Oscillatory shear rheology was used to determine the sol – gel transition temperature and the viscoelastic properties of S-GC-PNIPAM. The thermal gelling properties of the polymer give the resulting hydrogels viscous and elastic characteristics at physiological temperature. Storage modulus describes the energy stored in the elastic structure and the loss modulus describes the viscous component of the hydrogel. The solution to gel phase transition was determined to be at 35 °C for the hydrogel, allowing the material to be injectable at body temperature (Figure S3). As the temperature increases, an increase in storage and loss moduli was detected as the gelation process occurs, showing an increase in the stiffness of the gel. The hydrogel showed more elastic properties as the storage modulus was consistently higher than the loss modulus.

Elemental analysis by EDS was performed to determine the sulfur content on the internal scaffold surface. EDS is a standard procedure for identifying and quantifying elemental compositions of sample areas consisting of a micron or less. When comparing the two polymers, S-GC-PNIPAM shows an increase in sulfur content of 4.25% mass compared to no sulfur detected in GC-PNIPAM (Table S1). After analyzing the scaffolds with EDS, the samples were coated with gold and palladium for SEM imaging to examine the internal morphology of the scaffolds after gelation. SEM revealed a consistent porous structure for GC-PNIPAM and S-GC-PNIPAM, demonstrating that the sulfonation process did not alter the polymer morphology (Figure S4). The porous structure of chitosan provides support in tissue engineering therapies as it acts as a scaffold to provide the necessary support to physically guide differentiation and proliferation of cells for tissue growth responses [33]. Chitosan has been widely examined for tissue scaffolding due to its hydrophilic structure retaining water as well as maintaining bioactive proteins such as growth factors [34].

### Sequential and sustained protein release

S-GC-PNIPAM was examined for the ability to sequentially release three proteins according to their binding affinities to heparin, and in a sustained manner, with an *in vitro* release test study. Literature shows IL-10 having a strong heparin-binding affinity and a comparable affinity to FGF-2 (IL-10: K_d_ = 54 nM [23]; FGF-2: K_d_ = 74 nM [35]), while FGF-2 has a much stronger affinity for heparin sulfate than VEGF (FGF-2: K_A_ = 1.68 × 10^7^ M^-1^; VEGF: K_A_ = 8.14 × 10^5^ M^-1^) [36]. This suggests that IL-10 has an increased binding affinity for the sulfonate groups on S-GC-PNIPAM compared to VEGF. Additionally, PDGF demonstrated an equilibrium binding constant (K_A_ = 1.33 × 10^6^ M^-1^ to heparin [36]) higher than VEGF, indicating it should bind stronger to S-GC-PNIPAM. According to these heparin binding affinities, VEGF should be released from the hydrogel first, followed by IL-10, with PDGF released last. The release profile for S-GC-PNIPAM shows that the sulfonate groups provide sustained and sequential release of the proteins (Figure 2).

**Figure 2.**
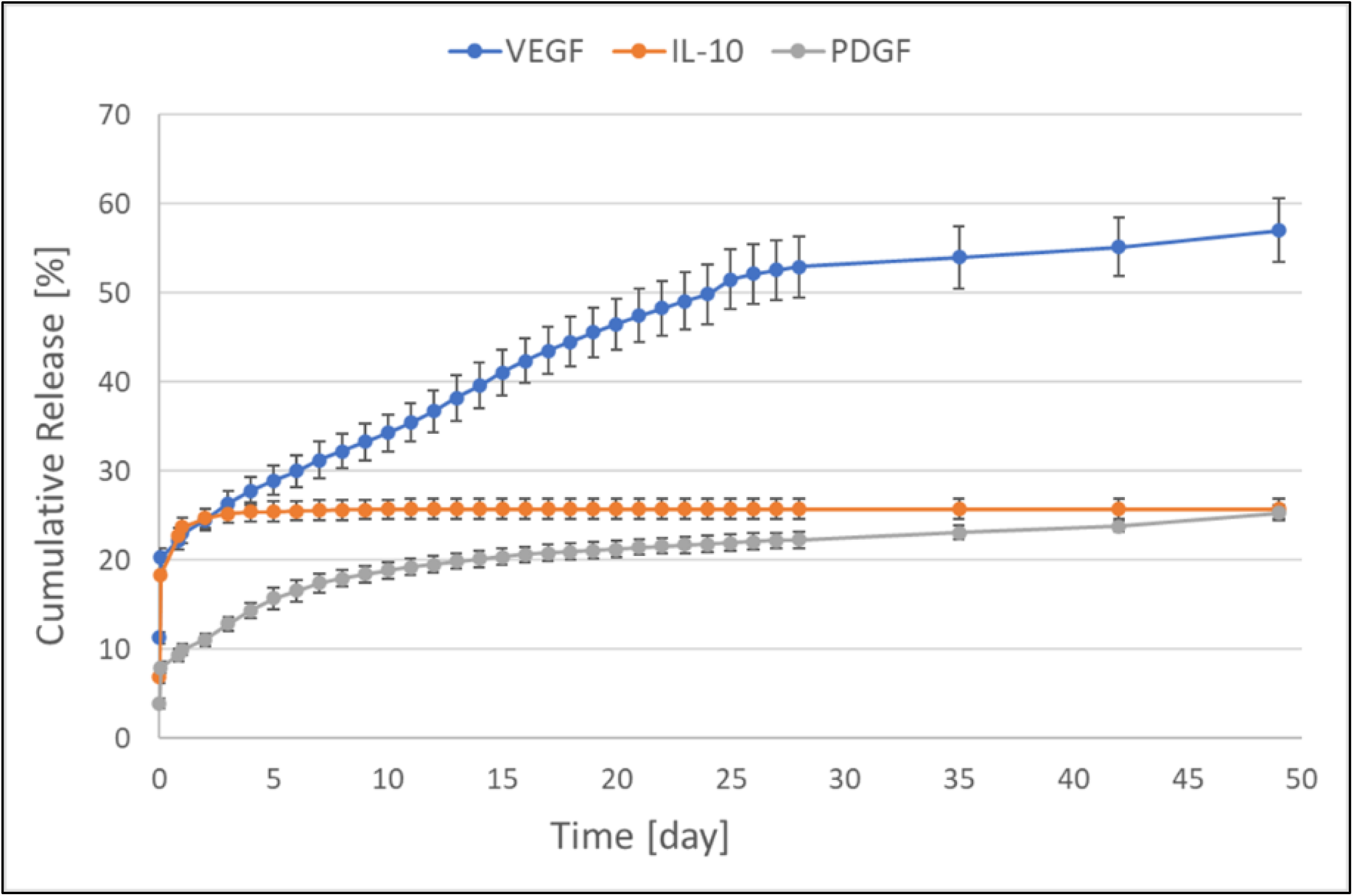
Cumulative release profile showing the sequential and sustained release of VEGF, IL-10 and PDGF from S-GC-PNIPAM. Error bars represent standard deviation.

The loading efficiencies were 89% for VEGF, 93% for IL-10, and 96% for PDGF, showing minimal burst release for all proteins. By day 3, 26% of loaded VEGF was released, with a sustained release rate of 0.93 ng per day through 49 days. IL-10 showed a reduced burst release compared to VEGF, suggesting a stronger electrostatic interaction to S-GC-PNIPAM with approximately 25% released by day 3. However, after day 3, only minute picogram amounts of IL-10 could be detected until it could not be detected by day 18. The lower cumulative release of IL-10 was likely due to its rapid degradation as recombinant human IL-10 has a short half-life of 2.7 to 4.5 h [22]. Additionally, the protein could be trapped within the polymer scaffold. PDGF exhibited the smallest burst release with approximately 13% of the protein released from the hydrogel by day 3. PDGF had a sustained release rate of 0.44 ng per day after day 0. By comparing the sustained release rates between VEGF and PDGF, it is apparent that the sulfonate groups on S-GC-PNIPAM are mimicking the protein binding function of heparin by reducing the release rate of PDGF by more than half. The different heparin-binding affinities seem to be responsible for sequentially releasing all three proteins, while providing a relatively linear release of VEGF and PDGF. Delivering these proteins in a sequential and sustained manner should promote long-term vascularization and reduce persistent inflammation in ischemic tissue, promoting optimal cardiac repair for treatment of MI.

### Improved cardiac function with S-GC-PNIPAM gel injections

The therapeutic benefits of spatiotemporally delivering VEGF, IL-10, and PDGF were evaluated in an acute MI mouse model. After MI, echocardiography can be utilized to measure cardiac function with ejection fraction, fractional shortening, and left ventricle inner diameter. These measurements were used to evaluate cardiac function after acute MI injury treated with intramyocardial injections of the thermal gel system loaded with the three proteins and using sham, saline, bolus three proteins, and S-GC-PNIPAM hydrogel alone as controls (Figure 3). Ejection fraction improved for all treatment groups compared to the saline control group (Figure 3A). Intramyocardial injections of S-GC-PNIPAM loaded with or without the proteins showed significant increases in ejection fraction compared to saline after 28 days (Figure 3A). The hydrogel injection groups appeared to have similar treatment effects for ejection fraction and fractional shortening. Potentially, S-GC-PNIPAM could increase left ventricular wall thickness by acting as a bulking material, thereby maintaining left ventricular geometry and improving cardiac function [37]. Although both thermoresponsive gel groups demonstrated statistically significant improvements compared to saline after 28 days, S-GC-PNIPAM loaded with the three proteins was the only group to show a significant improvement in ejection fraction and fractional shortening compared to the bolus three proteins group after 7 days (Figure 3A, B). The sequential delivery group exhibited a significant increase in fractional shortening to the saline group, after 28 days, while the other treatments groups did not (Figure 3B). Additionally, controlled delivery group demonstrated the smallest left ventricular inner diameter compared to the controls (Figure 3C). This shows evidence that the spatiotemporal delivery of the proteins is providing additional therapeutic benefits to cardiac function beyond the favorable mechanical properties of S-GC-PNIPAM itself.

**Figure 3.**
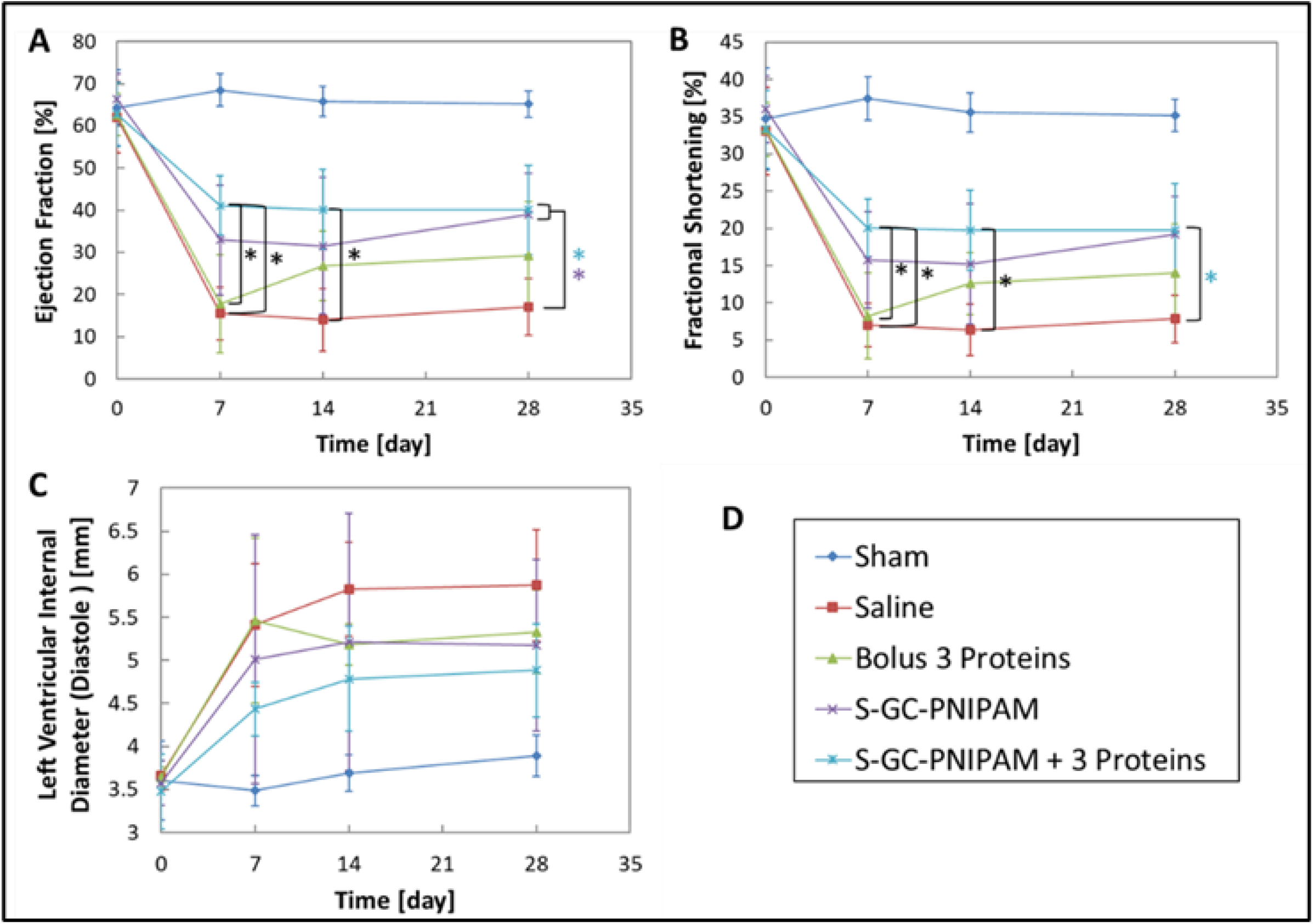
Echocardiography measurements of cardiac function after MI and intramyocardial injections. (A) ejection fraction, (B) fractional shortening, (c) left ventricular internal diameter at diastole. Data are presented as means and error bars represent standard deviation. * indicates *p* < 0.05.

Infarct size can be used to evaluate the potential treatment effect of intramyocardial injections on cardiac repair after MI. Masson trichrome staining identifies healthy myocardium that appears in red and pink from the infarct areas that are characteristic of fibrotic or collagenous tissue formation that appears in blue (Figure S5A). Infarct size was quantified by several measurement techniques, including area, endocardium and epicardium length, and midline length [38]. While there were no significant differences between the injection groups, S-GC-PNIPAM + 3 proteins exhibited the smallest infarct size for all measurement techniques (Figure S5B).

### Controlled delivery of therapeutic proteins from S-GC-PNIPAM promotes long-term vascularization in infarcted cardiac tissue

The restoration of blood flow to the ischemic heart through therapeutic angiogenesis is critical to tissue regeneration and functional protection of the myocardium. To investigate this angiogenic response, immunohistochemistry was performed to identify and quantify the different cell types involved in the process of blood vessel formation 28 days after MI. Endothelial cells were stained with CD31, and VWF was used to identify functional vascular endothelial cells and vessels when endothelial cells were stained with both VWF and CD31 (Figure 4A). Pericyte cell (mural cells or vascular smooth muscle cells) recruitment was observed by staining with α-SMA to identify long-term vascularization with co-staining of CD31 (Figure 4B). The quantification of immunohistochemical data for vascularization shows an increase in functional vascular endothelial cell vessels and mature vessels with pericyte recruitment after intramyocardial injections of S-GC-PNIPAM loaded with the three proteins (Figure 4C, D). The sequential delivery hydrogel group showed a significant increase in functional vascularization compared to the saline, bolus proteins, and S-GC-PNIPAM injection control groups (Figure 4C). The bolus injection of free proteins showed a higher number of VWF+ vessels compared to saline and the hydrogel alone, but there were no significant differences between the groups. In contrast, the significant increase in vascularization from the controlled delivery of the proteins from the hydrogel compared to the bolus protein injection indicates that the sequential release of VEGF and PDGF improved the formation of functional neovessels.

**Figure 4.**
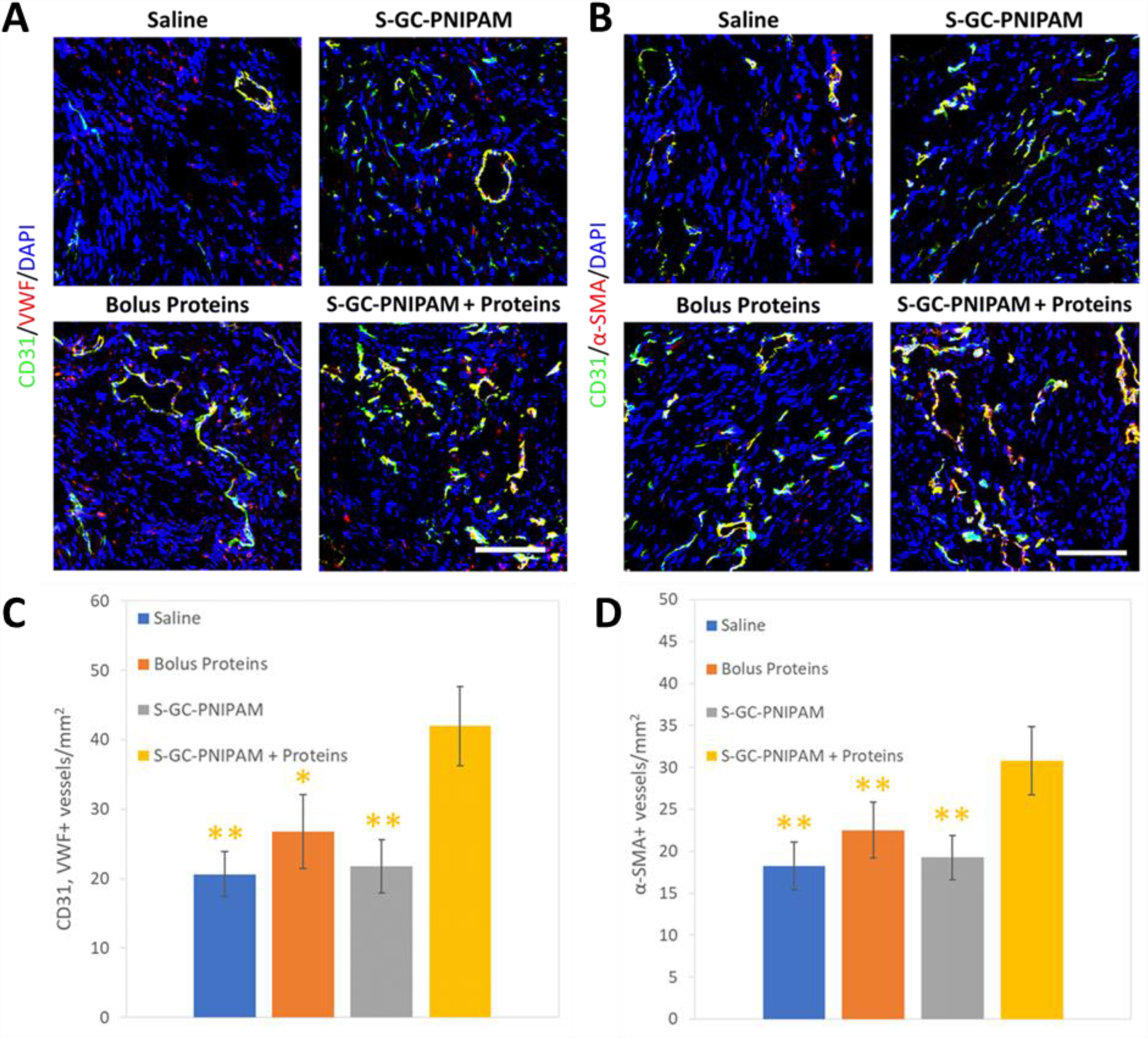
Immunohistochemical assessment of functional and mature vascularization by vessel formation 28 days after intramyocardial injections in acute MI mouse model. (A) Representative images show co-staining of CD31 (green), VWF (red) and DAPI (blue) Functional vascular cells were characteristic of CD31+ and VWF+ stains. (B) Representative images show co-staining of CD31 (green), α-SMA (red) and DAPI (blue). Mature vessels were characteristic of α-SMA+ with CD31+ associated stains. The scale bar represents 100 μm. (C) Quantification of vessel counts by functional vascular endothelial cells. (D) Quantification of vessel counts by pericyte association to endothelial cells. Data are presented as means and error bars represent standard deviation. * indicates *p* < 0.05, ** indicates *p* < 0.01.

As newly formed vessels can become hemorrhagic and regress over time if they are not stabilized with pericytes, it is important to promote mature and stable vasculature formation with therapeutic angiogenesis for beneficial cardiac repair. Few α-SMA+ vessels were observed in the saline, bolus proteins, and S-GC-PNIPAM injection groups with no statistical difference between them. However, the sequential delivery hydrogel group showed a significant increase in pericyte recruitment to the vessels compared to all control groups (Figure 4D). These results demonstrate the importance of releasing PDGF after the delivery of VEGF and in a sustained manner. This strong angiogenesis response is likely contributing to the observed improvement in cardiac function compared to the controls.

### Inflammatory response following intramyocardial thermoresponsive gel injections in acute MI mouse model

While recent studies on developing therapies for cardiac repair have mainly focused on revascularization strategies, limiting the over-reactive and extended inflammation response following MI is an important goal towards improving recovery of the myocardium [39]. To evaluate the local inflammatory response, immunohistochemistry was performed to identify and quantify macrophage infiltration by staining with CD68 (Figure 5A). In addition to examining the effect of IL-10 delivery on reducing inflammation in the heart, the biocompatibility of S-GC-PNIPAM can be determined by comparing the relative response seen after MI with the saline control injection.

**Figure 5.**
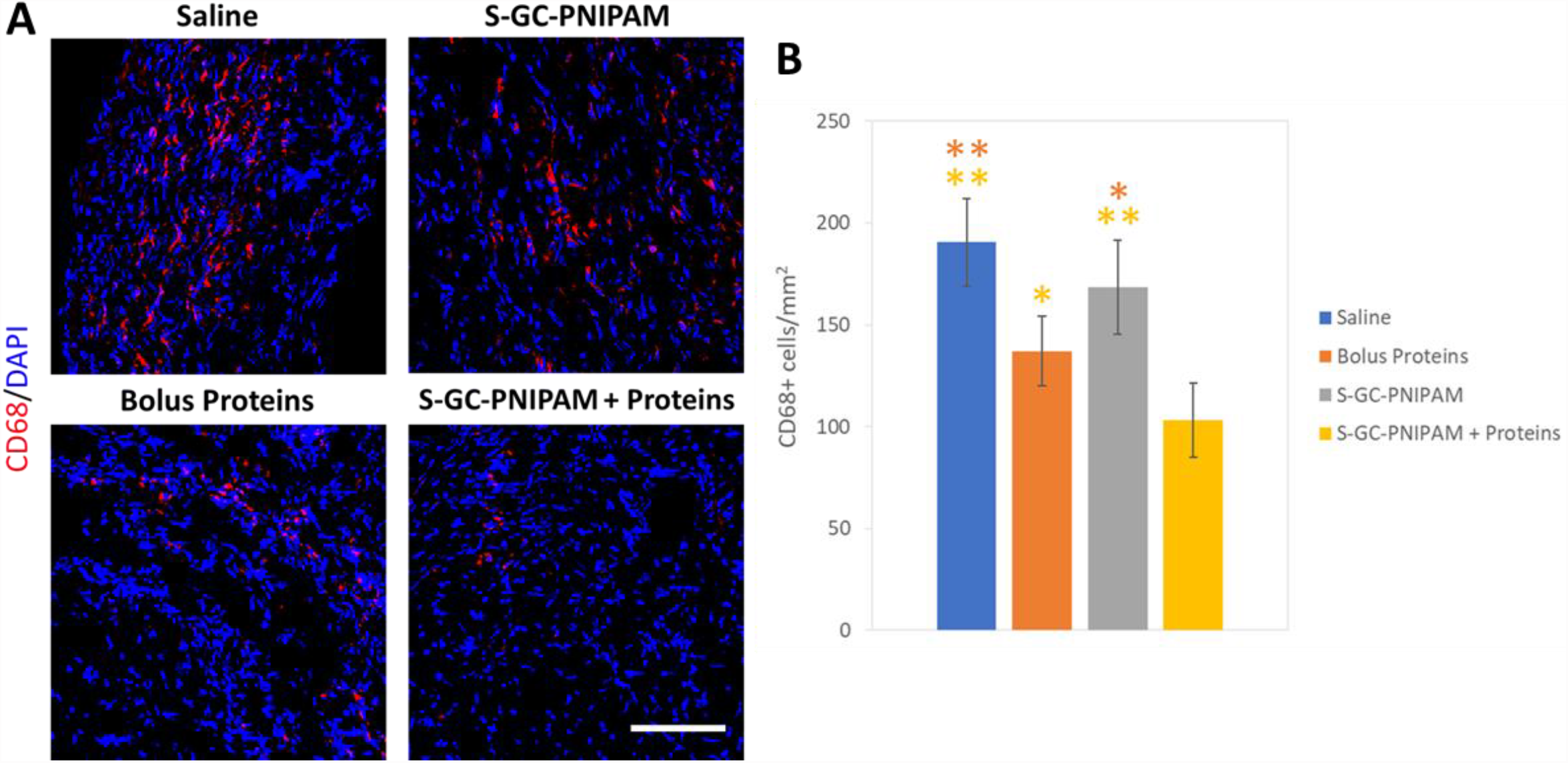
Immunohistochemical assessment of inflammatory response by macrophage infiltration 28 days after intramyocardial injections in acute MI mouse model. (A) Representative images show staining of CD68 (red) and DAPI (blue). Scale bar represents 100 μm. (B) Quantification of inflammatory response by macrophage cell count. Data are presented as means and error bars represent standard deviation. * indicates *p* < 0.05, ** indicates *p* < 0.01.

S-GC-PNIPAM loaded with the three proteins significantly reduced macrophage infiltration compared to the saline, bolus proteins, and S-GC-PNIPAM injection controls (Figure 5B). No statistical difference in the number of macrophages was observed between the hydrogel alone and the saline group demonstrating the biocompatibility of S-GC-PNIPAM and a minimal foreign body response (Figure 5B). The bolus proteins injection showed significantly reduced macrophages compared to saline and hydrogel injections confirming results seen from previous studies on IL-10 injections after MI [19,21]. These results suggest that the sustained delivery of IL-10 from the scaffold increases its long-term immunoregulatory effectiveness and potentially inhibits chronic inflammation post-MI.

### Factorial design of experiments study reveals optimal doses and combinations of proteins to increase vascularization and reduce inflammation

The focus of this work was on developing a new protein delivery system to protect cardiac function following MI; however, the cardiac function data for S-GC-PNIPAM with and without the proteins appears similar. To better optimize the delivery system, a design of experiments (DOE) study was utilized to determine the optimal protein doses for inducing therapeutic angiogenesis and limiting inflammation. Implementing a fractional factorial DOE study provides statistical models on different parameters while using a limited number of animals. The design of this controlled release study is based on the data from the *in vitro* release profile and acute MI mouse model study with additional input from literature on beneficial therapeutic proteins involved in different phases of healing post-MI [15,40–44]. S-GC-PNIPAM has demonstrated capabilities to deliver positively charged proteins according to their heparin-binding affinities with a sustained release over seven weeks. This should allow for the delivery of other proteins in a similar manner in relation to their affinity to sulfonate groups. To further elucidate other protein interactions involved in angiogenesis and cardiac repair, FGF-2 was included as a fourth protein for the factorial DOE study. FGF-2 has a high heparin-binding affinity, comparable to IL-10 affinity but a stronger affinity compared to VEGF, which should allow it to release in a sustained fashion similar to these proteins [36].

For this study, statistical software was used to generate a table of eight experimental dosage groups with different upper and lower doses for each protein (Table 1). A subcutaneous injection mouse model was used for this study since previous successful results have been produced using this model for evaluating other injectable gel systems for controlled protein delivery [32,45,46]. The main efficacy outcomes measured were functional vascularization, mature vascularization, and inflammation. Immunohistochemistry was used to quantify the different cell types involved in the process of blood vessel formation and inflammation with stains for CD31 and VWF to identify functional, vascular endothelial cells, CD31 and α-SMA for mature vasculature, and CD68 for macrophages. Based on previous studies, a sample size was determined as 3 mice per dose group. Vascularization should be significantly different in the subcutaneous region of the mice after injecting pro-angiogenic growth factors in a controlled fashion compared to the saline control. Therefore, to produce the statistical models, the minimum of 3 mice is sufficient.

The results from the immunohistochemical analysis showed a significant increase in neovascularization for dosage groups 1 and 4 compared to saline and all other injection groups (Figure 6A, Figure S6A). The six other experimental groups showed an increasing trend in vascularization compared to saline but with no statistical differences, likely due to increased variability (Figure S6A). Additionally, dosage groups 1, 2, and 4 all showed a significant increase in functional vascular endothelial cells compared to saline (Figure 7A). The analysis of variance modeling results demonstrated significant main effects on promoting functional vascularization from VEGF and FGF-2 (*p* < 0.001), with negligible effects from IL-10 (*p* = 0.177) and PDGF (*p* = 0.655) (Table S2). VEGF and FGF account for 95% of the total sum of squares for the model and appear to dominate the main effect of increasing vascular endothelial cells. VEGF had the greatest main effect on increasing vascularization, accounting for 62% of the total sum of squares, followed by FGF accounting for 33% (Table S2). The significant positive slopes on the main effects plot for VEGF and FGF-2 indicates that increasing vascularization is best achieved by injecting the higher doses of the two proteins (Figure 9). As VEGF and FGF-2 are two of the most significant regulators of angiogenesis, the combination of these proteins in a controlled delivery format provides beneficial effects on the early stages of vessel formation.

**Figure 6.**
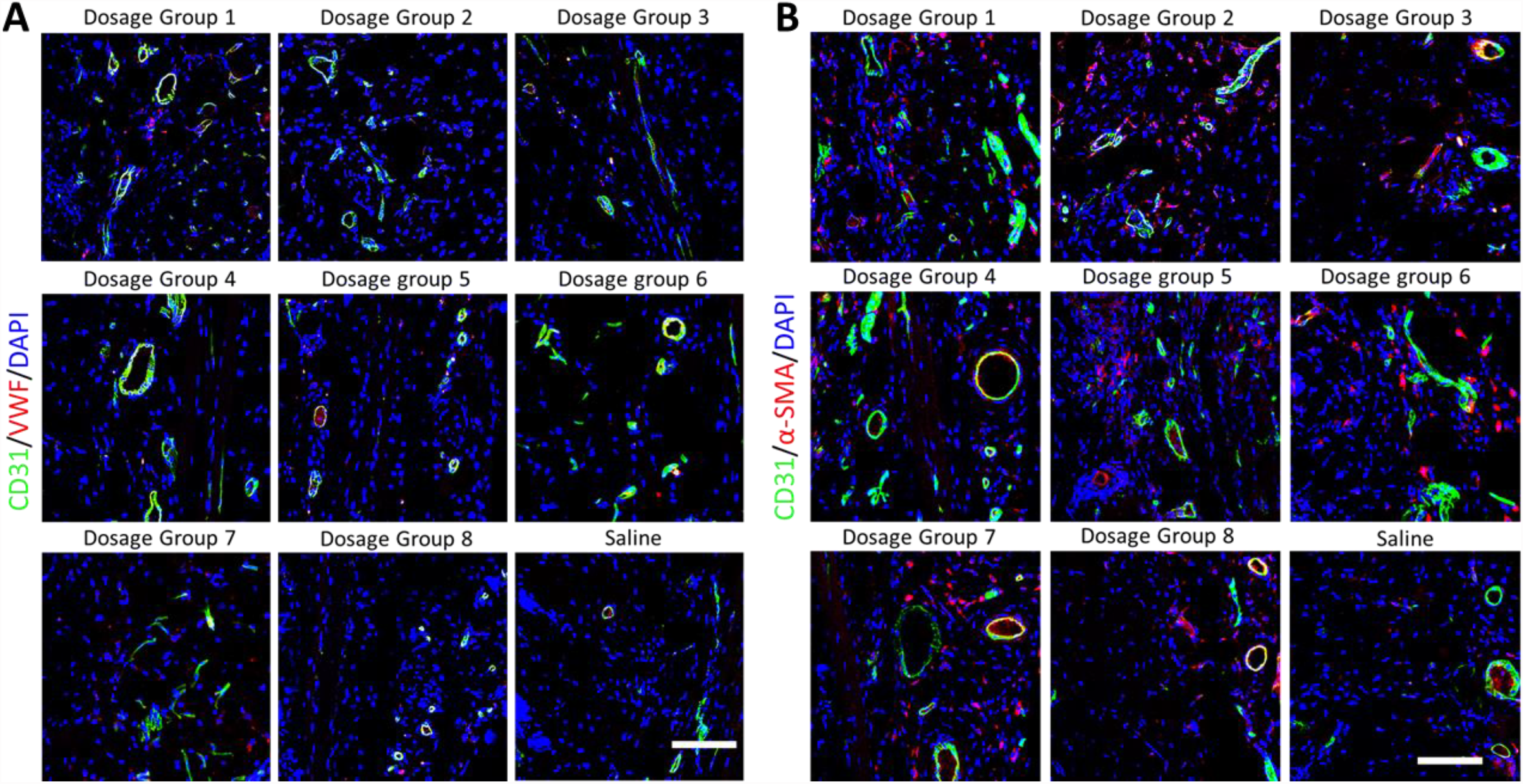
Immunohistochemical assessment of functional and mature vascularization 28 days after subcutaneous injections of dose groups. (A) Representative images show co-staining of CD31 (green), VWF (red) and DAPI (blue). Functional vascular cells were characteristic of VWF+ with CD31+ associated stains. (B) Representative images show co-staining of CD31 (green), α-SMA (red) and DAPI (blue). Mature vessels were characteristic of α-SMA+ with CD31+ associated stains. The scale bar represents 100 μm.

**Figure 7.**
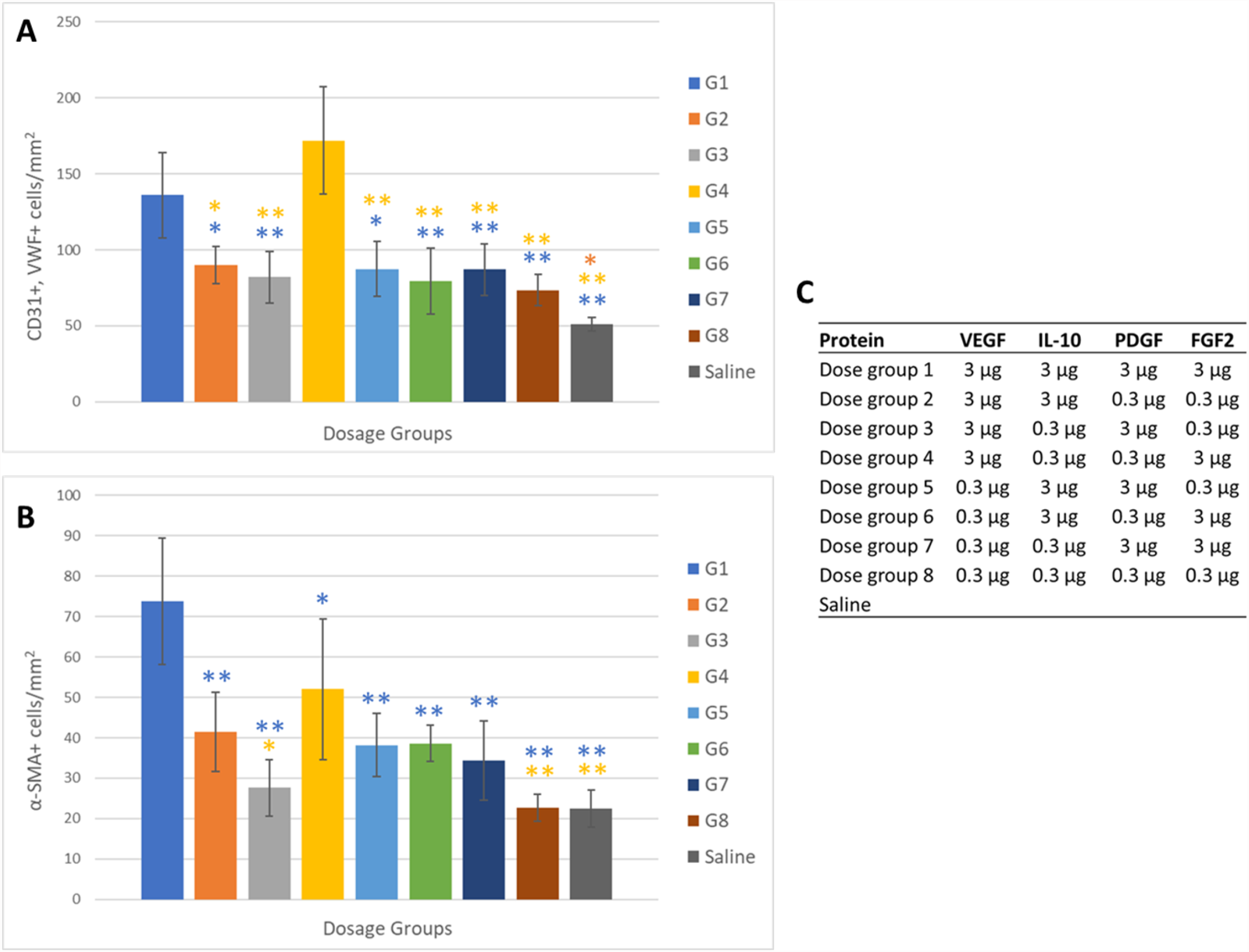
Quantification of immunohistochemical assessment functional and mature vascularization 28 days after subcutaneous injections of dose groups. (A) Cell counts by functional vascular endothelial cells. (B) Cell counts by pericyte association to endothelial cells. (C) Table showing the dosages for each group. G1 from the legend represents dose group 1 in the table and each other group is represented, respectively. Data are presented as means and error bars represent standard deviation. * indicates *p* < 0.05, ** indicates *p* < 0.01.

The results from the immunohistochemical analysis on mature vascularization showed a significant increase in CD31+ endothelial cells for all protein loaded hydrogel groups compared to saline (Figure 6B, Figure S6B). Dosage group 1 promoted significantly more endothelial cells compared to all other hydrogel groups as well (Figure S6B). For long-term vascularization, dosage groups 1 and 4 both showed significantly higher pericyte cell density compared with the saline control, dosage group 3, and dosage group 8 (Figure 7B). Additionally, dosage group 1 recruited significantly more α-SMA+ cells compared to all other groups. The analysis of variance modeling results demonstrated significant main effects on promoting stable vascularization from VEGF, IL-10, and FGF-2 (*p* < 0.001), with less noticeable effects from PDGF (*p* = 0.152) (Table S2).

VEGF and FGF account for 70% of the total sum of squares for the model and displayed the greatest main effects on inducing long-term vascularization. FGF-2 showed the largest main effect on pericyte recruitment, accounting for 39% of the total sum of squares, followed by VEGF accounting for 31%, then IL-10 with 26% (Table S2). The significant positive slopes on the main effects plot for VEGF, IL-10, and FGF-2 indicates that increasing stable vessel formation is optimally attained by injecting the higher doses of the three proteins (Figure 9). Interestingly, the difference in dosage for PDGF appears to have a minor effect on recruiting pericytes to the newly formed vessels. It seems that a low dose of PDGF released from the hydrogel, in combination with high doses of the other proteins, provides sufficient benefits for promoting stable angiogenesis. As FGF-2 induces the migration of vascular smooth muscle cells similar to PDGF, these results indicate that the higher dose of FGF-2 is providing a more significant cellular recruitment signal compared to a high dose of PDGF.

The results from the inflammatory response assessment showed a significant decrease in macrophage infiltration for all hydrogel groups loaded with a high dose of IL-10 compared to all injection groups with a low dose of IL-10 except for dosage group 8 (Figure 8A, B). However, dosage groups 1 and 2 showed a significant reduction in CD68+ cells in comparison to dosage group 8 as well. Dosage groups 1, 2 and 5, all with the high dose of IL-10, showed no statistical difference in inflammatory cell presence to the saline control group (Figure 8B). The analysis of variance modeling results exhibited significant main effects on decreasing inflammation from only IL-10 (*p* < 0.001), with a small effect from FGF-2 (*p* = 0.057), and negligible effects from VEGF and PDGF (*p* = 0.658, *p* = 0.685) (Table S2). IL-10 displayed a major effect on reducing the inflammatory response from the foreign material as it accounts for 85% of the total sum of squares for the model. The significant negative slope on the main effects plot for IL-10 shows that lowering the inflammatory response is best accomplished by injecting the higher dose of the protein (Figure 9).

**Figure 8.**
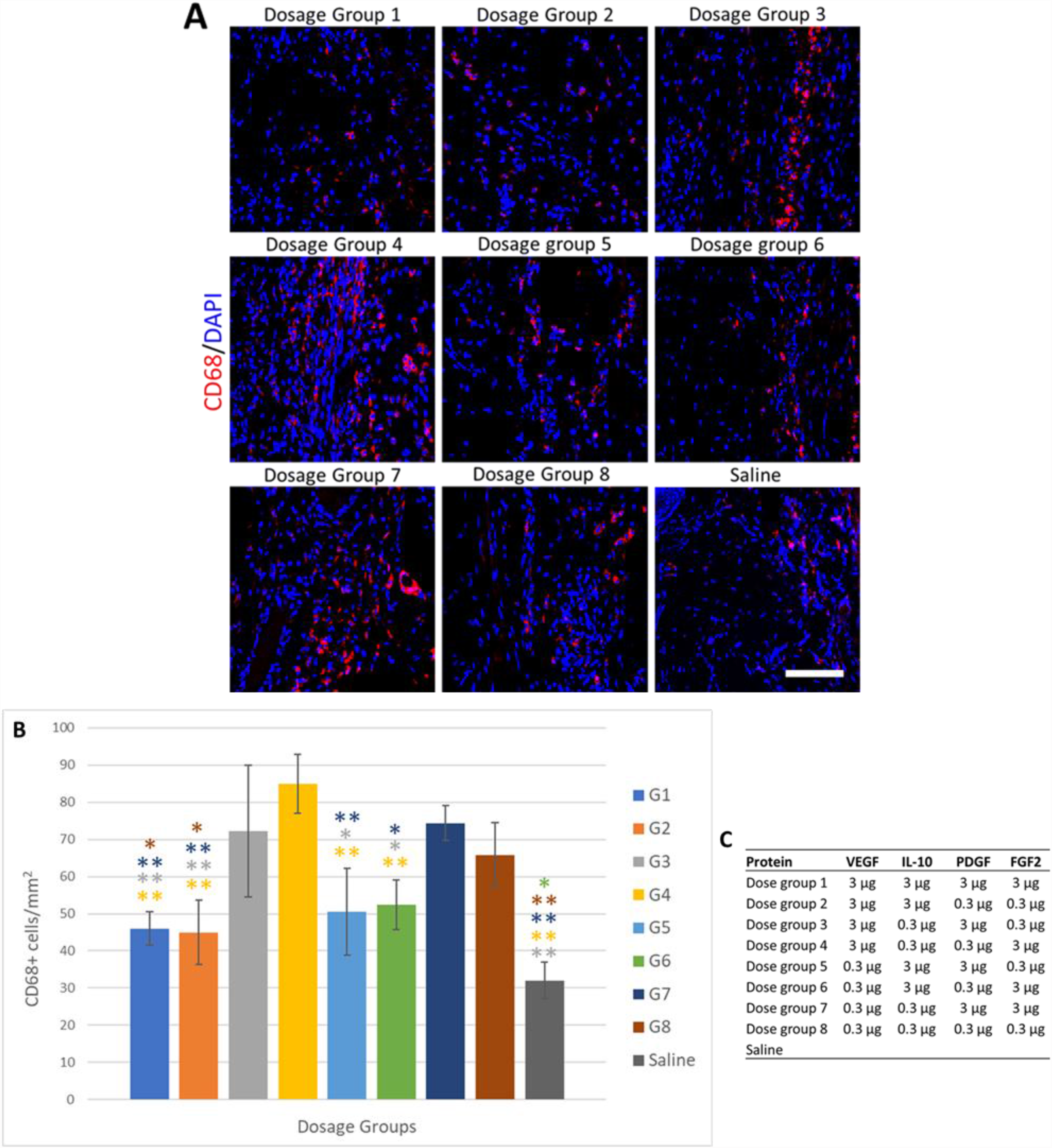
Immunohistochemical assessment of inflammatory response by macrophage cell count 28 days after subcutaneous injections of dose groups. (A) Representative images show staining of CD68 (red) and DAPI (blue). Scale bar represents 100 μm. (B) Quantification of macrophages by cell count. (C) Table showing the dosages for each group. G1 from the legend represents dose group 1 in the table and each other group is represented, respectively. Data are presented as means and error bars represent standard deviation. * indicates *p* < 0.05, ** indicates *p* < 0.01.

**Figure 9.**
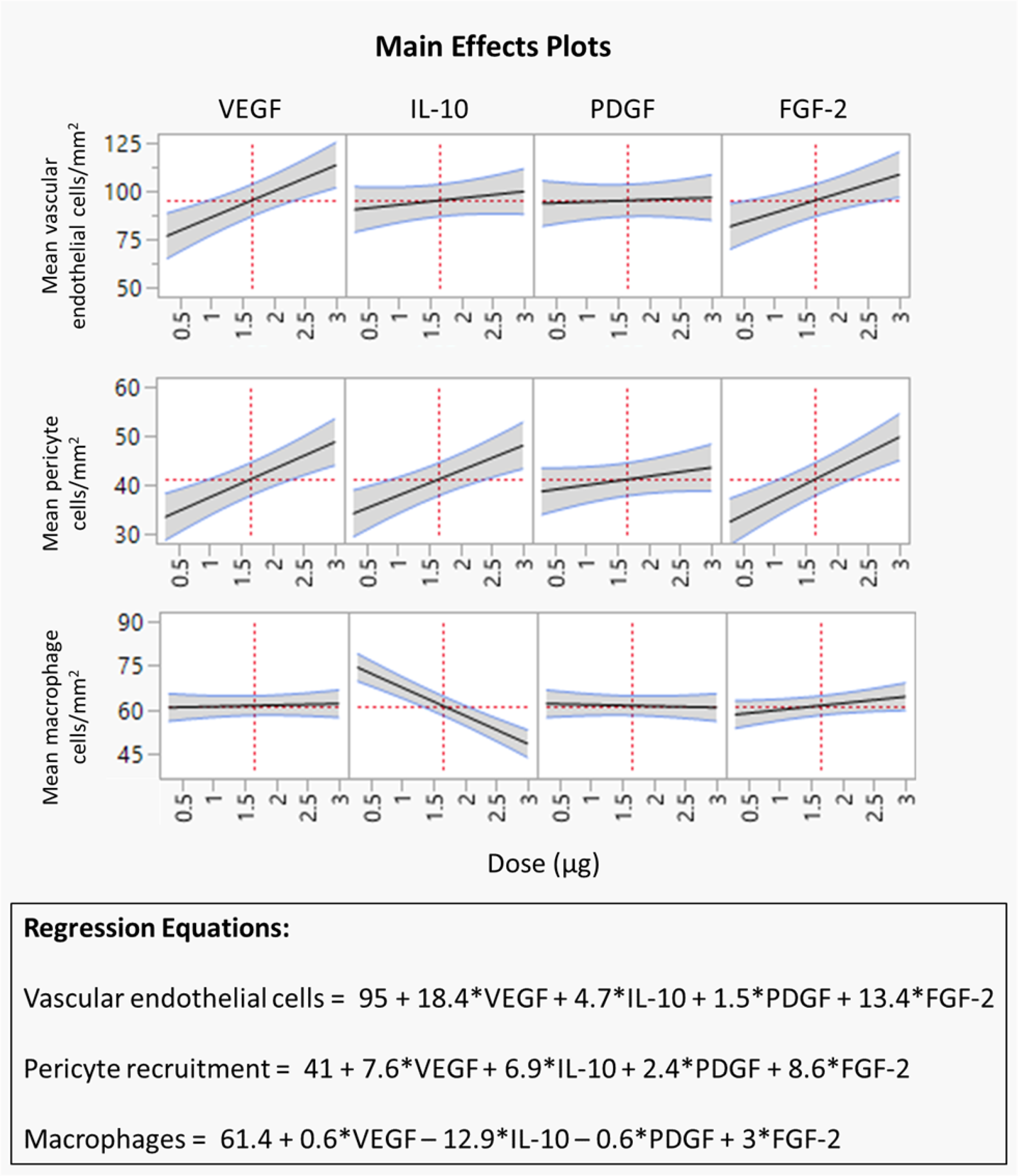
Main effects plots and regression model equations shows the individual effect of each protein on functional vascularization, pericyte recruitment, and macrophage infiltration from the lower and upper doses.

## DISCUSSION

MI results in progressive cardiomyocyte cell death and harmful pathological remodeling that can culminate in heart failure. Interventional therapies utilized in the acute phase of healing following MI can reduce the progression of these events and promote cardiac repair. While minimally invasive cell therapies have shown promise for preventing adverse remodeling, this approach is limited by inadequate cardiomyocyte cell sources and poor survival and engraftment rates [47,48]. Injectable biomaterials can overcome the limitations of cell-based therapies as they can be delivered minimally invasively to induce endogenous repair mechanisms. This strategy aims to reduce left ventricular wall stress by mechanical load shielding and increasing ventricle wall thickness, which moderates adverse cardiac remodeling [49]. Many natural biological materials, including chitosan, gelatin, fibrin, and alginate, have been examined as injectable materials for myocardial infarction treatments. These materials have been used in a variety of tissue engineering applications due to their biocompatible properties and being readily available for investigators. Many preclinical acute MI rodent studies have demonstrated improved ventricular remodeling and cardiac function by intramyocardially injecting biomaterials that can act as tissue bulking agents [50–52]. In this study, an injectable sulfonated thermoresponsive gel, S-GC-PNIPAM, was developed and investigated for localized and controlled protein delivery to protect cardiac function following MI. We showed that intramyocardial injections of the hydrogel significantly improved ejection fraction in an acute MI mouse model, suggesting that the hydrogel alone can prevent harmful ventricular remodeling. To enhance the reparative benefits seen from injecting our hydrogel, we examined the controlled delivery of therapeutic proteins from the polymer scaffold.

After a myocardial infarction, the adult heart has minimal regeneration potential. However, protein therapies have the potential for inducing tissue regeneration in the damaged myocardium. Many protein candidates have been identified to have cardiac regeneration capabilities including growth factors to stimulate angiogenesis, chemotactic factors to attract stem cells, and other proteins that promote cardiomyocyte mitogenesis and stem cell growth and differentiation [53]. As some approaches to angiogenesis involve the use of viral vectors or cells, protein delivery is the most simple and direct way [46]. VEGF and FGF-2 are well-studied growth factors and have been tested in human clinical trials. These clinical studies used direct injection of free proteins and failed to demonstrate efficacy, likely due to their rapid diffusion rate, poor biostability, and short half-lives in vivo [54]. Controlled delivery systems for therapeutic proteins are highly desirable and currently under extensive research as these systems deliver proteins directly to the treatment site and can stabilize the biologics [46,55,56]. However, there remains significant barriers to protein delivery systems related to loading capacity and long-term efficacy [15]. We have shown that S-GC-PNIPAM can overcome these limitations by evaluating the delivery system with an *in vitro* release study. The hydrogel demonstrated a very high loading efficiency for VEGF, IL-10 and PDGF with minimal burst release and a sustained release of the proteins over seven weeks. Sulfonating the polymer allowed for the sequential release of the proteins according to their binding affinities to heparin.

We investigated the benefits of controlled release of VEGF, IL-10 and PDGF from S-GC-PNIPAM using an acute MI mouse model. The pilot study was performed to examine the potential therapeutic effects of S-GC-PNPAM loaded with the proteins by evaluating cardiac function, quantifying vascularization, and measuring the inflammatory response. Intramyocardial injections of the hydrogel loaded with the three proteins was compared to sham, saline, bolus three proteins, and the polymer alone as control groups. Treatment with S-GC-PNIPAM + proteins was the only group to show significant improvement in cardiac function for ejection fraction and fractional shortening compared to saline and bolus proteins after seven days. The controlled delivery group demonstrated significant increases in ejection fraction and fractional shortening compared to saline after 28 days as well. The bolus proteins group did not show significant improvements in cardiac function compared to saline, demonstrating the importance of spatiotemporal release of the proteins from S-GC-PNIPAM. The smallest infarct size was observed after injection of the hydrogel with the proteins, while significant increases in functional and stable vascularization were seen compared to all other groups. The controlled delivery of IL-10 significantly decreased macrophage infiltration post-MI, and the biocompatibility of S-GC-PNIPAM was demonstrated. These results suggest that the hydrogel delivery system can promote robust angiogenesis and provide cardioprotective properties for the treatment of MI.

The cardiac function data did not show any significant differences for S-GC-PNIPAM with and without the proteins. This led us to optimizing the delivery system using a DOE study to determine optimal proteins doses and combinations for inducing therapeutic angiogenesis and decreasing inflammation in a subcutaneous injection mouse model. To our knowledge, this is the most comprehensive study to date on examining optimal protein doses for increasing long-term vascularization and reducing inflammation. VEGF, IL-10, PDGF, and FGF-2 all have comparatively specific roles they play in angiogenesis and cardiac repair after MI. VEGF and IL-10 are therapeutically advantageous early after MI to limit inflammation and induce angiogenesis, while PDGF needs to be delivered sequentially and in a sustained aspect to stabilize newly formed vessels. FGF-2 should be delivered in a sustained manner as it initiates angiogenesis, induces the migration of smooth muscle cells, and promotes the survival of cardiomyocytes [14]. All four of these proteins have demonstrated anti-apoptotic properties on cardiomyocytes as well [57–59]. The combination of these four proteins has never been tested in a controlled delivery format before. Thus, it is unknown how each protein will interact together in the aspects of angiogenesis and reducing inflammation, and for their potential in cardiac repair. This data could provide insights into the best combinations of proteins and dosages for use in future MI animal model studies by detecting different protein interactions that are important for designing these potential treatments.

The results from the DOE study showed that the most effective hydrogel injection group was dosage group 1, with high doses (3 μg) of each protein providing the largest increase in functional and stable angiogenic vessels and the most significant reduction in inflammatory cells from the foreign body response. Similar results were observed for the high and low doses of PDGF, signifying that future pro-angiogenic factor delivery studies could reduce the amount of this protein compared to the other biologics, reducing associated costs, or remove the protein entirely when using this combination of factors. Several studies have shown the importance of sequentially delivering PDGF after VEGF, and our results appear to conflict with current literature [13,17,41,60]. There are a few potential reasons why PDGF had an insignificant effect on promoting mature vascularization with the recruitment of pericytes. Greenberg et al. demonstrated that releasing VEGF and PDGF simultaneously results in complete suppression of angiogenesis in a chorioallantoic membrane model and subcutaneous injection mouse model [16]. The high dose of PDGF in our study, in combination with the other proteins, could be overloading the capacity of S-GC-PNIPAM, causing a burst release of PDGF, and disabling the sequential delivery component of our delivery system. We are planning to perform a release study with all four of these proteins for a future manuscript to determine if this is the case.

As this combination of proteins has not been evaluated before, the pro-angiogenic effects of IL-10 and FGF-2 could have a more significant effect on recruiting pericytes and vascular smooth muscle cells compared to PDGF. Our data shows that delivering a higher dose of IL-10 exhibited a significant effect on promoting mature vascularization. This data supports results from a previous controlled release study where Chen et al. observed an increase in vascular smooth muscle cell density when increasing the dose of IL-10 while keeping the amount of FGF-2 the same in an acute MI mouse model [61]. M2c macrophages are activated by IL-10 and studies have shown that these regulatory macrophages promote angiogenesis *in vivo* and secrete high levels of MMP9, a protease associated with vascular remodeling [62,63]. As FGF-2 stimulates the proliferation and migration of vascular smooth muscle cells, this growth factor could be working in tandem with IL-10 to promote vascular remodeling and mature vascularization, rendering the delivery of PDGF negligible. Additionally, Kano et al. exhibited that synergism between VEGF and FGF-2 enhance intercellular PDGF signaling with VEGF increasing endothelial cell PDGF expression and FGF-2 increasing pericyte PDGF receptor β expression [64]. This study also showed that exogenous PDGF decreased pericyte recruitment when used in combination with VEGF and FGF-2 in an in vivo Matrigel plug assay. These studies and the data presented here indicate that the combination of VEGF, IL-10, and FGF-2 in high doses may have more significant effects on inducing long-term and stable vascularization compared to combinatorial protein delivery with PDGF.

The results from this study may guide investigators on selecting the best combination and doses of VEGF, IL-10, PDGF, and FGF-2 for future MI studies to optimally promote stable angiogenesis and limit inflammation in order to protect cardiac function. This factorial design approach may represent a compelling method for optimizing delivery systems as therapies for complex diseases. It allows for determining significant effects of each drug and drug interactions, while reducing associated costs by examining half of the dose combinations compared to a full factorial study. There are some limitations to this study: (1) The acute MI study was performed with a small number of animals which reduces statistical power. (2) The MI study endpoint was only four weeks, potentially not showing long-term benefits from the controlled protein delivery compared to the hydrogel alone as it begins to degrade. (3) A subcutaneous injection model was used for the DOE study, and this tissue does not have the same mechanical properties as the heart. However, this preclinical animal model has been used by many investigators to evaluate angiogenic protein delivery before proceeding into more clinically relevant MI animal models where they have seen successful results translate from one model to the other [46,64–66].

Several experiments will be designed to address these issues and subsequently published in a future manuscript. We will increase the animal numbers for our next MI study to ensure the related evaluations are sufficiently powered. The optimal protein doses discovered from the DOE analysis will be evaluated in this future MI study, and one experimental injection group will not include PDGF to determine if the combination of VEGF, IL-10, and PDGF provides increased benefits compared to all four together. Evaluating this optimal dose group with the MI model will determine the effectiveness of the DOE subcutaneous injection model. S-GC-PNIPAM + proteins showed significant improvements in cardiac function compared to the saline control, and this optimized injectable system could provide additional benefits towards ejection fraction compared to the hydrogel group. An additional timepoint of eight weeks will be used to determine the long-term effects of the controlled delivery of the proteins from S-GC-PNIPAM. The increase in functional and mature vessel formation and decrease in macrophages suggests that there should be longer lasting effect from the sequential protein delivery on protecting cardiac function. This data may provide valuable information for improving future studies on controlled protein delivery for MI treatments by determining the best combinations of proteins to promote cardiac repair.

## CONCLUSIONS

We developed a novel multiple protein delivery system that was synthesized, characterized, and evaluated for its therapeutic potential with *in vivo* animal studies. The molecular structure of S-GC-PNIPAM was confirmed through ^1^H NMR, FTIR and elemental analysis. An *in vitro* release study demonstrated the controlled delivery capabilities of the delivery system by sequentially releasing VEGF, IL-10 and PDGF in a sustained manner over seven weeks. These results showed that the sulfonate groups on S-GC-PNIPAM are electrostatically binding to the positively charged proteins, mimicking the growth factor binding properties of heparin and preserving their function. The protection of cardiac function following intramyocardial thermoresponsive gel injection in an acute MI model was investigated. Treatment with S-GC-PNIPAM + proteins showed significant improvement in cardiac function for ejection fraction and fractional shortening, while increasing long-term vascularization and decreasing inflammation compared to controls. A DOE subcutaneous injection model showed that the delivery of higher doses of VEGF, IL-10, and FGF-2 from S-GC-PNIPAM provided enhanced blood vessel growth and reduced macrophage infiltration, while PDGF had no significant effects. These results demonstrate that S-GC-PNIPAM provides cardioprotective properties for the treatment of MI and can deliver therapeutic proteins in a spatiotemporal manner for regenerative medicine and tissue engineering applications.

## Funding

This work was supported by NIH/NCATS Colorado CTSA Grant Number UL1 TR002535, the National Institutes of Health T32 HL072738 and the American Heart Association AHA/17GRNT33661024.Contents are the authors’ sole responsibility and do not necessarily represent official NIH views.

## Notes

### Competing Interest Statement

The authors have declared no competing interest.

