## Supplemental Materials for "A Sulfonated Thermoresponsive Injectable Gel for Sequential Release of Therapeutic Proteins to Protect Cardiac Function After a Myocardial Infarction"

### Supplemental Figures and Tables

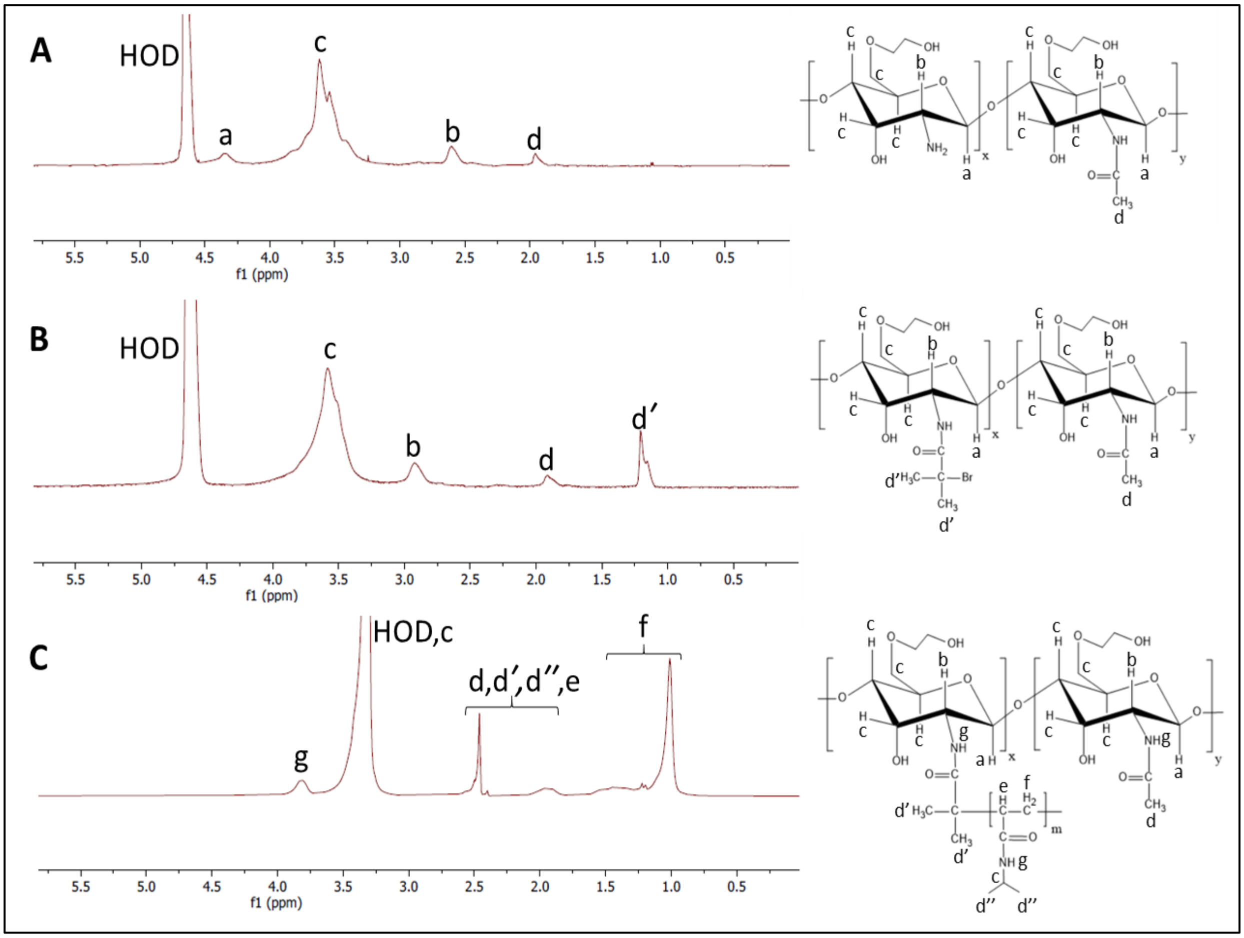


**Figure S1. $^1$H NMR spectra of (A) GC, (B) GC-Br, (C) GC-PNIPAM.** $^1$H NMR confirmed the successful conjugation of BMPA and the ATRP synthesis of PNIPAM onto GC.

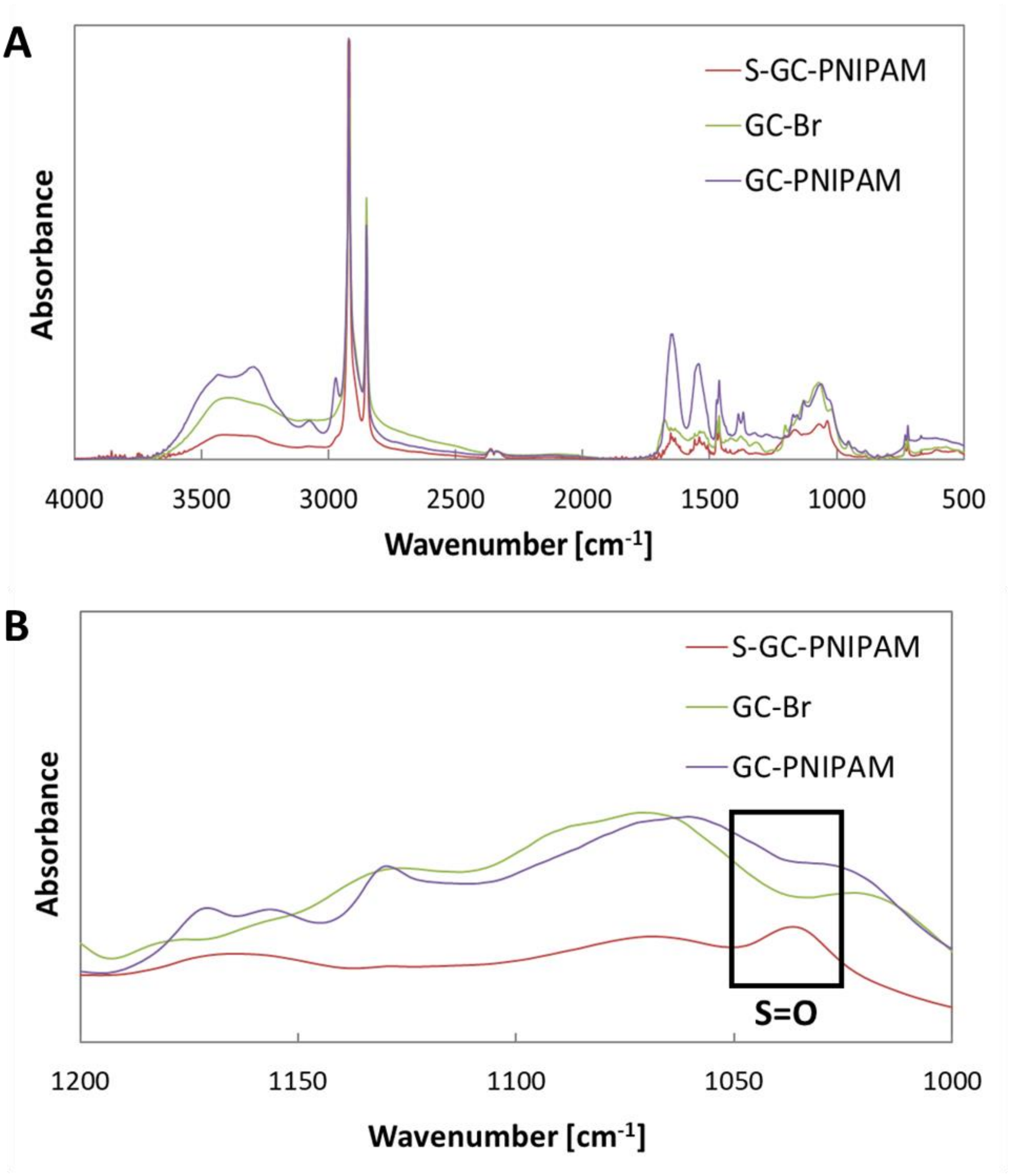


**Figure S2. FTIR spectrum confirming molecular structure and sulfonation of S-GC-PNIPAM.** (A) Full spectrum of GC-Br , GC-PNIPAM and S-GC-PNIPAM. (B) Spectrum highlighting specific wavenumber range of sulfonation peak and confirming sulfonation of S-GC-PNIPAM.

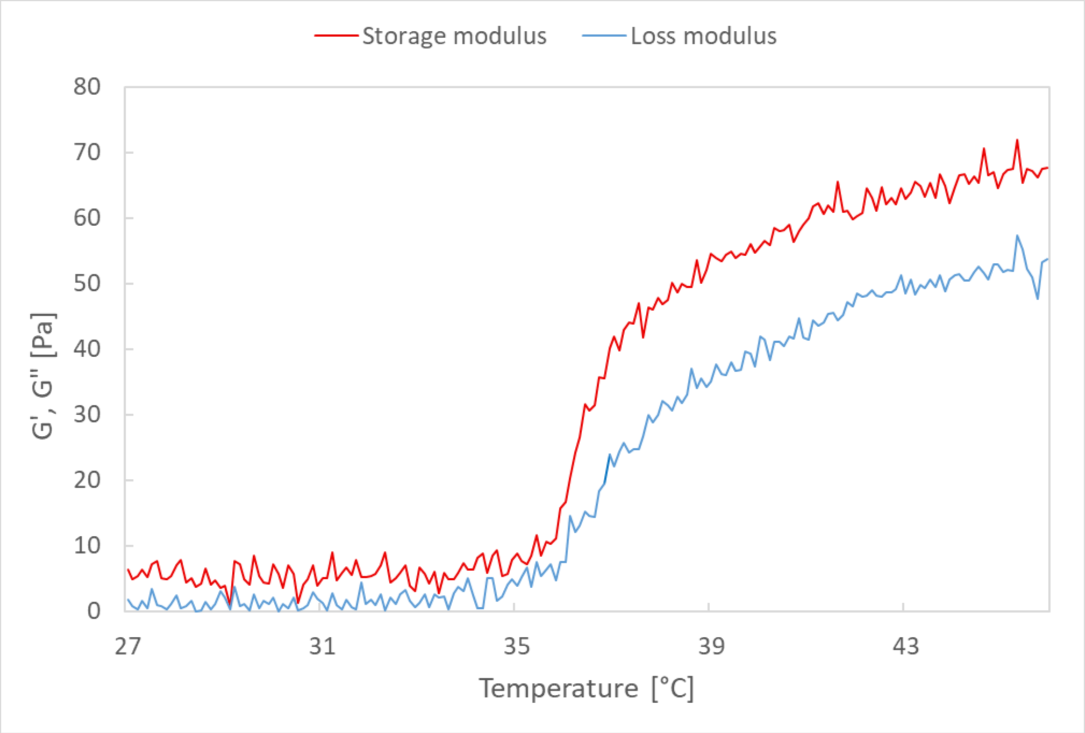


**Figure S3. Rheological behavior of S-GC-PNIPAM.** Measurements were taken at a frequency of 1 Hz and stress of 0.05 Pa. At temperatures above 34 ℃, the storage (G’) and loss (G”) moduli begin to increase rapidly demonstrating the liquid to gel phase transition.

| Scaffold | Oxygen [%] | Nitrogen [%] | Sulfur [%] | Sodium [%] | Chloride [%] |
|---|---|---|---|---|---|
| GC-PNIPAM | 35.54 | 16.64 | 0.00 | 20.07 | 27.75 |
| S-GC-PNIPAM | 46.80 | 15.93 | 4.25 | 13.68 | 19.34 |

**Table S1. Elemental analysis by EDS confirming the sulfonation of S-GC-PNIPAM.** Measurements were taken as mass percentages on the inner scaffold surface.

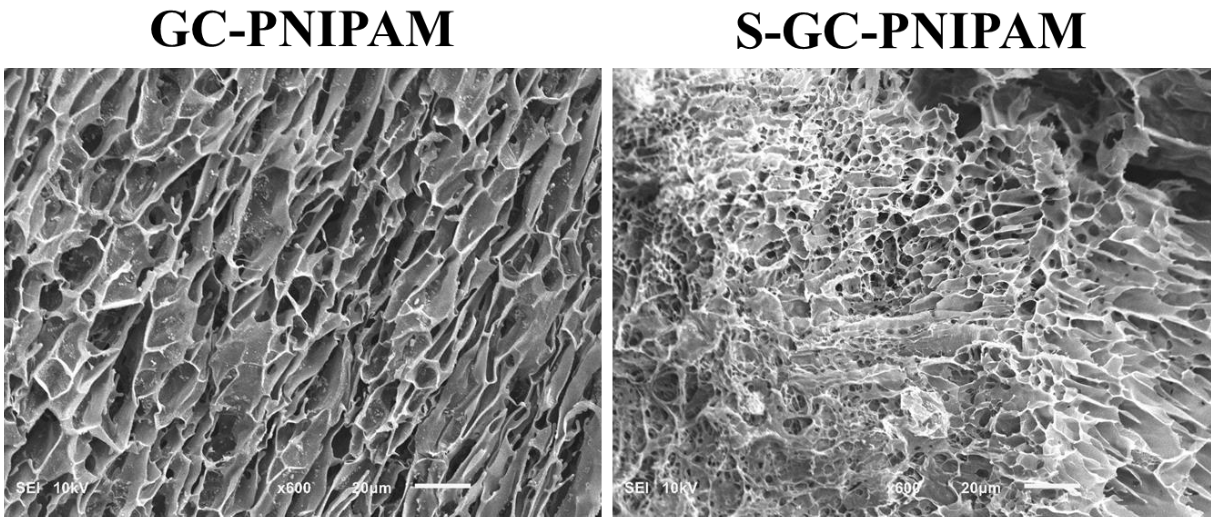


**Figure S4. SEM images of hydrogel scaffolds.** The images show the porous internal morphology of the hydrogels before and after sulfonation. Scale bar represents 20 µm.

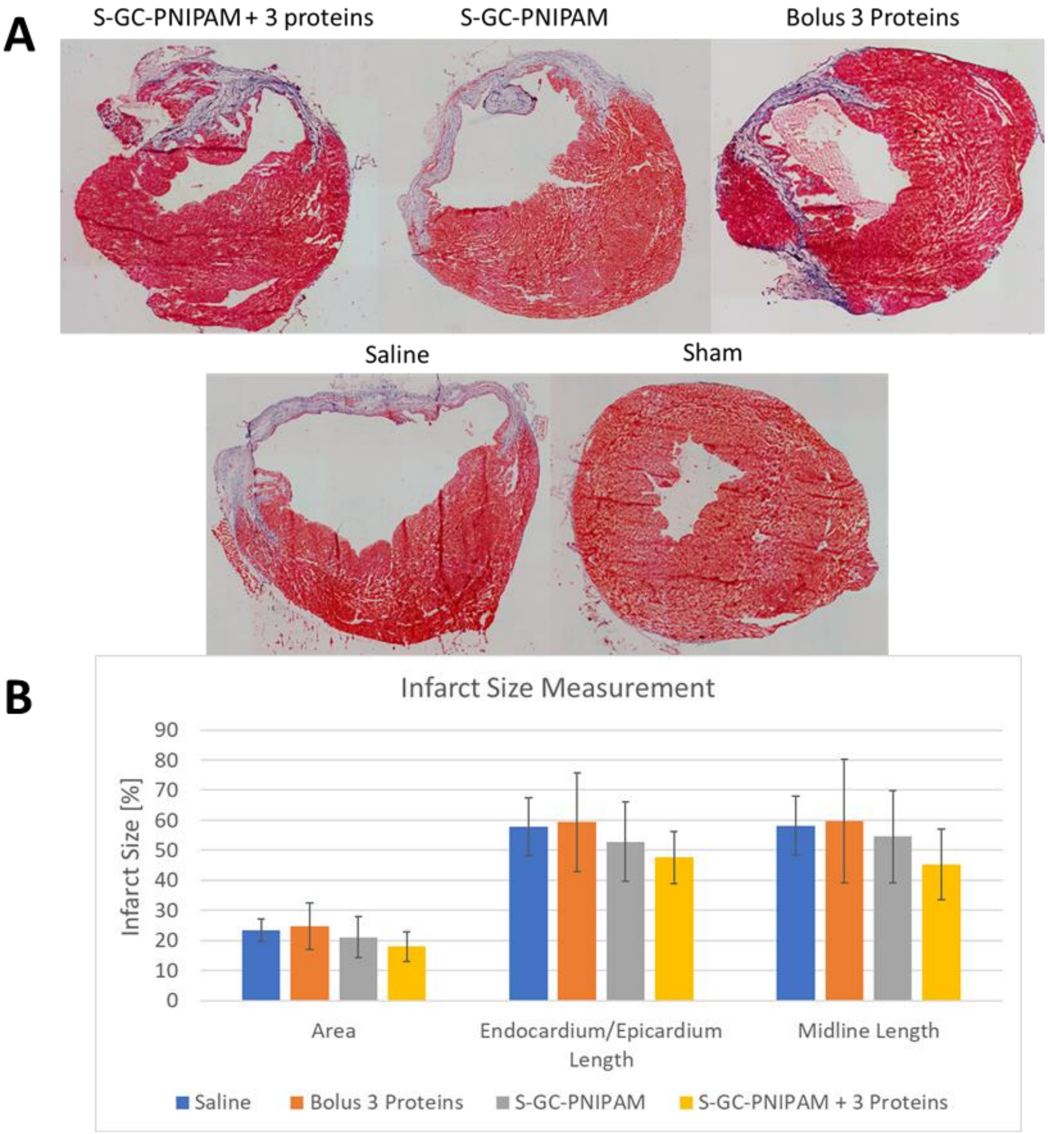


**Figure S5. Masson's trichrome stains of hearts and infarct size measurements.** Staining (A) and infarct size measurements (B) performed following acute MI, 28 days after permanent ligation. (A) Red staining shows muscle fibers, pink staining shows cytoplasm and blue staining shows fibrotic tissue. (B) Infarct size determined by area, endocardium/epicardium length and midline length measurements. Scale bar represents 1000 μm. Data are presented as means and error bars represent standard deviation.

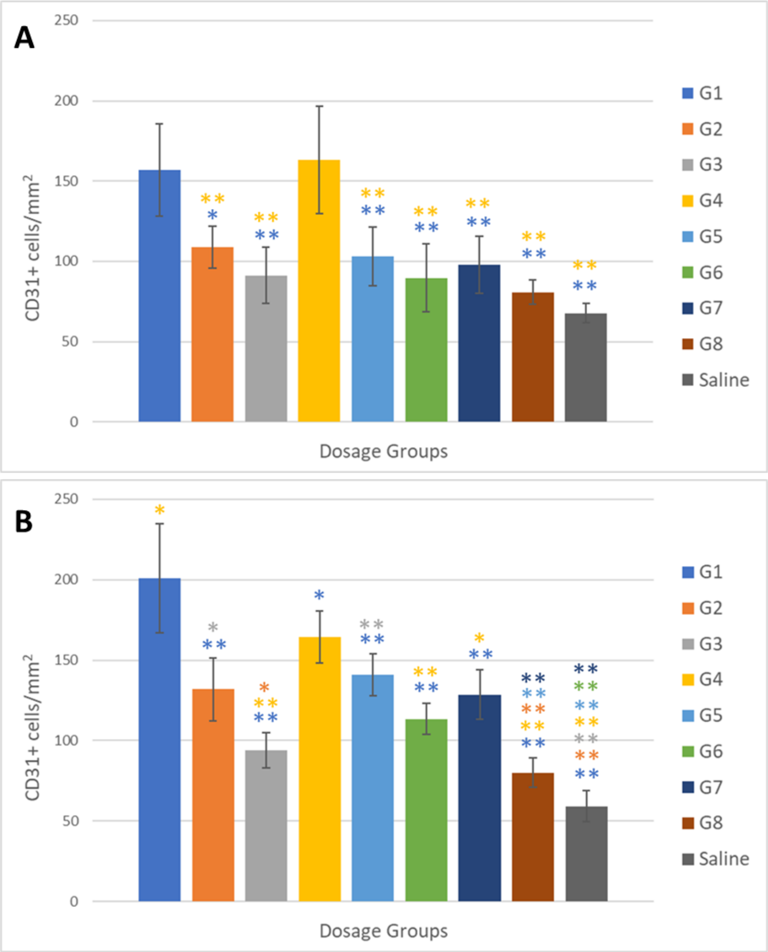


C

| Protein | VEGF | IL-10 | PDGF | FGF2 |
|---|---|---|---|---|
| Dose group 1 | 3 µg | 3 µg | 3 µg | 3 µg |
| Dose group 2 | 3 µg | 3 µg | 0.3 µg | 0.3 µg |
| Dose group 3 | 3 µg | 0.3 µg | 3 µg | 0.3 µg |
| Dose group 4 | 3 µg | 0.3 µg | 0.3 µg | 3 µg |
| Dose group 5 | 0.3 µg | 3 µg | 3 µg | 0.3 µg |
| Dose group 6 | 0.3 µg | 3 µg | 0.3 µg | 3 µg |
| Dose group 7 | 0.3 µg | 0.3 µg | 3 µg | 3 µg |
| Dose group 8 | 0.3 µg | 0.3 µg | 0.3 µg | 0.3 µg |
| Saline | | | | |

**Figure S6. Quantification of immunohistochemical assessment of neovascularization 28 days after subcutaneous injections of dose groups.** (A) Cell counts by endothelial cells co-stained with VWF. (B) Cell counts by endothelial cells co-stained with α-SMA. (C) Table showing the dosages for each group. G1 from the legend represents dose group 1 in the table and each other group is represented, respectively. Data are presented as means and error bars represent standard deviation. * indicates $p < 0.05$, ** indicates $p < 0.01$.

| Model | Sum of Squares | F-Ratio | P-Value |
|---|---|---|---|
| Functional Vascularization | | | |
| VEGF | 13505.6 | 29.5 | < 0.001 |
| IL-10 | 874.2 | 1.9 | 0.177 |
| PDGF | 93 | 0.2 | 0.655 |
| FGF-2 | 7209.2 | 15.7 | < 0.001 |
| Mature Vascularization | | | |
| VEGF | 2325.6 | 21.2 | < 0.001 |
| IL-10 | 1918.2 | 17.5 | < 0.001 |
| PDGF | 235.2 | 2.1 | 0.152 |
| FGF-2 | 2975.6 | 27.1 | < 0.001 |
| Inflammation | | | |
| VEGF | 15.6 | 0.2 | 0.685 |
| IL-10 | 6682.2 | 71.4 | < 0.001 |
| PDGF | 15.6 | 0.2 | 0.685 |
| FGF-2 | 366.0 | 3.9 | 0.057 |

**Table S2. Analysis of variance data**. Analysis of variance results show relative significance of each of the 4 proteins on promoting functional vascularization and pericyte recruitment and reducing macrophage infiltration.
